## Supplementary material for "Identification of microbial exopolymer producers in sandy and muddy intertidal sediments by compound-specific isotope analysis": Latex_zip_file_V2: ajout.tex

\begin{figure*}
\centering
\includegraphics[scale=0.7]{figures/Fig4.pdf}
\caption{C and N isotopic ratio of fatty acids and EPS fractions. \textbf{a}: $\delta^{13}$C of fatty acid biomarkers (corrected according to equation \ref{eq1}). \textbf{b}: $\delta^{13}$C and $\delta^{15}$N biplot of bound and colloidal EPS.
}\label{figure4}
\end{figure*}

\subsection*{C:N ratio of colloidal and bound EPS}\label{res:C:N_ratio}

The C:N ratios were related to the carbon isotope ratio (figure \ref{figure5}). Changes in $\delta^{13}$C values matched with shifts in the elemental content of the two EPS fractions. Bound EPS showed lower $\delta^{13}$C values (-21.08 $\pm$ 2.79\textperthousand) than colloidal EPS (-13.28 $\pm$ 3.59\textperthousand) and this difference was significant (Student t test: $t=-9.31, df=57, p=\num{4.9e-13}$ ). The$\ ^{13}$C depleted values corresponded to the higher C:N ratios. The relation between C:N ratio and $\delta^{13}$C corresponded to an exponential decrease (figure \ref{figure5}, $y=\exp(-x)$) with $y=\text{C:N ratio}$ and $x=\delta^{13}$C ($R^2=0.9913, p<\num{2.2e-16}$)


\begin{figure}
\centering
\includegraphics[scale=0.5]{figures/Fig5.pdf}
\caption{relation between C:N ratio and $\delta^{13}$C of colloidal and bound EPS. Muddy and sandy sites were grouped together. The black line corresponds to a non linear regression fit ($y=\exp(-x), R^2=0.9913, p<\num{2.2e-16}$). The grey area represents the confidence interval for a type I error $\alpha=0.999$.
}\label{figure5}
\end{figure}


% MBP isotopes
% mudflats la rochelle : d15N = 6 à 10, d13C = -13 à -17 (20:5w3 = -20 / décalage d'~5)  %https://www.sciencedirect.com/science/article/abs/pii/S0141113618304744
% Chez euglènes 18:3w3 et 14:0 = major FA (%https://www.jlr.org/content/5/3/352.long).
%De plus, 18:3w3 abondant en conditions mixotrophes et 14:0 abondant en condition hétérotrophe : %https://www.ncbi.nlm.nih.gov/pmc/articles/PMC4890288/
% Diatoms : The most common fatty acids are myristic acid (14:0), palmitic acid (C16:0), palmitoleic acid (C16:1n-7), DHA, and EPA (Opute, 1974; Jiang et al., 2016). %https://www.frontiersin.org/articles/10.3389/fmars.2017.00017/full#:~:text=Most%20fatty%20acids%20in%20diatoms,et%20al.%2C%202016).
% Fungi : rich in 18:1n-9 & 18:2n-6 (https://www.mdpi.com/1660-3397/18/8/379/htm). # 22:5w6 possible signature %fungi https://books.google.fr/books? id=xYg8AAAAIAAJ&pg=PA92&lpg=PA92&dq=18:2w6+euglena&source=bl&ots=ZlpeJW8ODY&sig=ACfU3U2IrzQlI6DateXcD9tNWOH8SP7NOQ&hl=fr&sa=X&ved=2ahUKEwj9nuC6-MLtAhVFJBoKHaeJBfwQ6AEwBHoECAkQAg#v=onepage&q=18%3A2w6%20euglena&f=false  
% Cyanobacteria : rich in 16:0, 16:1∆9 (16:1∆9 16:1ω7 16:1n-7), 18:1∆9 ou 18:1n-9 %(file:///Users/cedric.hubas/Downloads/Maslova2004_Article_LipidFattyAcidCompositionAndTh.pdf) file:///Users/cedric.hubas/Downloads/Oren1985_Article_UnsaturatedFattyAcidCompositio.pdf 
% Cyano : %file:///Users/cedric.hubas/Downloads/Wada-Murata1998_Chapter_MembraneLipidsInCyanobacteria.pdf
% nomenclature : %https://www.economie.gouv.fr/files/files/directions_services/dgccrf/Scl/Publication/2015%20Publi%20SCL13%20-%20Analyse%20des%20lipides%20-%20tehcniques%20de%20l'ing%C3%A9nieur%20-%20Extraction-param%C3%A8tres%20physico-chimiques%20-%20constituants%20majeurs.pdf
% Besides well-known bacterial ability to adapt FA synthesis under environmental conditions, it was demonstrated that bacteria can incorporate free exogenous FA from its environment to its membrane (%https://hal.archives-ouvertes.fr/hal-01608805/document). 
% It was recently shown that S. aureus can incorporate exogenous FA via a fatty acid kinase-dependent pathway (Parsons et al., 2014). . SFA and UFA diffuse passively through the cytoplasmic membrane and bind, respectively, to FakB1 and FakB2 fatty acid binding proteins before they are converted to acyl carrier protein. In biofilm cells, as in planktonic cells, SFA and UFA can be incorporated and elongated. 


\section*{Discussion}

As emphasised by Hirave et al. \cite{Hirave2020UnderstandingCSIA} degradation does not impact isotopic ratio of long-chain fatty acids and n-alkanes in comparison to bulk $\delta^{13}$C analysis.


Main conclusions of the paper are that EPS production and degradation are supported by very different communities in muddy and sandy sediments and that EPS sources are more diverse in the sand. In the mud, bound EPS come mainly from diatoms and the colloidal EPS are the result of a degradation of these EPS (by certain specialised bacteria). In sand it is more diverse. We find the same pattern as in the mud with a contribution of diatoms but there is also a production of bound EPS which is possibly of bacterial origin (or cyanobacteria ?). Also, fatty acids indicate that the bacteria that degrade EPS are not the same as those that produce EPS.


Relation between d13C and C:N ratio indicated that shifts in d13C values are linked to shifts in EPS precursor material \cite{Hood2005SourcesIsotopes} 

Multiple studies of litter decomposition have demonstrated that litter δ15N increases as decay proceeds. https://link.springer.com/article/10.1007/s11104-015-2542-1
une diminution du C/N indique une reminéralisation de la MO par les microorgansimes. Yes decomposition very much dependent upon C:N ratio. Wider the C:N ratio lesser the decomposition rate of the material.  %https://en.wikipedia.org/wiki/Carbon-to-nitrogen_ratio#:~:text=When%20composting%2C%20microbial%20activity%20utilizes,of%2020%2D30%3A1.


% SUPP TABLE
\begin{table}[]
\caption{Presumed sources of colloidal and bound EPS (carbohydrates, proteins) at muddy and sandy sites. \textbf{Bold} = major fatty acid (20-40\%) in the corresponding sources. No specific font-type  = important fatty acid (10-20\%), \textit{Italic} = present in trace amounts (<10\%)}
\label{tab:table2}
\begin{tabular}{@{}lllll@{}}
\toprule
Location & EPS type & Fatty acids & Position based on d13C & Possible origin of FA \\ \midrule
\multicolumn{1}{c}{\multirow{15}{*}{Mud}} & \multirow{4}{*}{Colloidal} & 22:6n-3 & \multirow{2}{*}{Aligned} & Haptophyta, Dinoflagelate (51), \\
\multicolumn{1}{c}{} &  & 20:0 &  & lorem (XX) \\ \cmidrule(l){3-5} 
\multicolumn{1}{c}{} &  & 18:4n-3 & \multirow{2}{*}{Overlying} & Haptophyta, Pheophyceae (49b), Diatoms (44) \\
\multicolumn{1}{c}{} &  & 17:0iso &  & Bacteria (52,53) \\ \cmidrule(l){2-5} 
\multicolumn{1}{c}{} & \multirow{11}{*}{Bound} & 20:4n-6 & \multirow{3}{*}{Aligned} & lorem (XX) \\
\multicolumn{1}{c}{} &  & 17:1n-5/7 &  & lorem (XX) \\
\multicolumn{1}{c}{} &  & 16:4n-1 &  & lorem (XX) \\ \cmidrule(l){3-5} 
\multicolumn{1}{c}{} &  & 22:0 & \multirow{8}{*}{Overlying} & lorem (XX) \\
\multicolumn{1}{c}{} &  & 20:5n-3 &  & lorem (XX) \\
\multicolumn{1}{c}{} &  & 18:4n-6 &  & lorem (XX) \\
\multicolumn{1}{c}{} &  & 18:2n-6 &  & lorem (XX) \\
\multicolumn{1}{c}{} &  & 18:1n-9/18:3n-3 &  & lorem (XX) \\
\multicolumn{1}{c}{} &  & 17:0, 17:iso &  & lorem (XX) \\
\multicolumn{1}{c}{} &  & 16:3n-4/16:4n-3 &  & lorem (XX) \\
\multicolumn{1}{c}{} &  & 15:0 &  & lorem (XX) \\ \midrule
\multirow{21}{*}{Sand} & \multirow{15}{*}{Colloidal} & 22:6n-3 & \multirow{13}{*}{Aligned} & lorem (XX) \\
 &  & 20:5n-3 &  & lorem (XX) \\
 &  & 20:4n-6 &  & lorem (XX) \\
 &  & 18:4n-7 &  & lorem (XX) \\
 &  & 18:1n-9/18:3n-3 &  & lorem (XX) \\
 &  & 18:1n-7 &  & lorem (XX) \\
 &  & 17:1n-5/7 &  & lorem (XX) \\
 &  & 17:0 &  & lorem (XX) \\
 &  & 16:3n-4/16:4n-3 &  & lorem (XX) \\
 &  & 16:1n-7 &  & lorem (XX) \\
 &  & 16:1n-5 &  & lorem (XX) \\
 &  & 16:0 &  & lorem (XX) \\
 &  & 15:0 &  & lorem (XX) \\ \cmidrule(l){3-5} 
 &  & 14:0 & \multirow{2}{*}{Overlying} & lorem (XX) \\
 &  & 18:0 &  & lorem (XX) \\ \cmidrule(l){2-5} 
 & \multirow{6}{*}{Bound} & 15:0iso, 15:0anteiso & Aligned & lorem (XX) \\ \cmidrule(l){3-5} 
 &  & 18:1n-7 & \multirow{5}{*}{Overlying} & lorem (XX) \\
 &  & 18:0 &  & lorem (XX) \\
 &  & 16:0 &  & lorem (XX) \\
 &  & 15:0 &  & lorem (XX) \\
 &  & 14:0 &  & lorem (XX) \\ \bottomrule
\end{tabular}
\end{table}
