## Supplementary material for "Identification of microbial exopolymer producers in sandy and muddy intertidal sediments by compound-specific isotope analysis": Latex_zip_file_V2: ArticleV1.0.tex

\documentclass[times, twocolumn]{zHenriquesLab-StyleBioRxiv}
%\documentclass[a4paper, 11pt]{article}
\usepackage{textcomp}
\usepackage[utf8]{inputenc}
\usepackage{booktabs}
\usepackage{multirow}
\usepackage{blindtext}
\usepackage{soul}
\usepackage{graphicx}
\usepackage{lineno}
\usepackage{lipsum}
\usepackage{siunitx}
\usepackage{ulem}
%\usepackage{draftwatermark}

% Different numbering for supplementary figures and tables
\newcommand{\beginsupplement}{%
        \setcounter{table}{0}
        \renewcommand{\thetable}{ST\arabic{table}}%
        \setcounter{figure}{0}
        \renewcommand{\thefigure}{SF\arabic{figure}}%
     }

\linenumbers
\leadauthor{Hubas} 

\begin{document}

\title{Identification of microbial exopolymer producers in sandy and muddy intertidal sediments by compound-specific isotope analysis.}
\shorttitle{Stable isotope ratios of EPS}

\author[1,$\ast$]{Cédric Hubas}
\author[1]{Julie Gaubert-Boussarie}
\author[2]{An-Sofie D’Hondt}
\author[3]{Bruno Jesus}
\author[4,5]{Dominique Lamy}
\author[3]{Vona Meleder}
\author[1,3]{Antoine Prins}
\author[3]{Philippe Rosa}
\author[2]{Willem Stock}
\author[2]{Koen Sabbe}

\affil[1]{Laboratoire de Biologie des Organismes et Ecosystèmes Aquatiques (BOREA) Muséum National d'Histoire Naturelle, CNRS, Sorbonne Université, IRD, Université des Antilles, Université de Caen Normandie; Station Marine de Concarneau, Place de la croix,29900, Concarneau, France}
\affil[2]{Department of Biology, Research Group Protistology and Aquatic Ecology, Ghent University, Krijgslaan 281/S8, 9000, Belgium}
\affil[3]{Nantes Université, Institut des Substances et Organismes de la Mer, ISOMer, UR 2160, F-44000 Nantes, France}
\affil[4]{Laboratoire de Biologie des Organismes et Ecosystèmes Aquatiques (BOREA) Muséum National d'Histoire Naturelle, CNRS, Sorbonne Université, IRD, Université des Antilles, Université de Caen Normandie; Jardin des plantes, Bâtiment arthropodes, Paris, France}
\affil[5]{Institute of Ecology and Environmental Sciences of Paris (iEES-Paris), Sorbonne Université, Univ Paris Est Créteil, IRD, CNRS, INRA, 4 place Jussieu, 75005 Paris, France}


\maketitle
%\SetWatermarkText{Preprint}
\begin{abstract}
Extracellular polymeric substances (EPS) refer to a wide variety of high molecular weight molecules secreted outside the cell membrane by biofilm microorganisms. In the present study, EPS from marine microphytobenthic biofilms were extracted and their isotope ratios were analysed. A comparison of these ratios with the carbon isotope ratios of fatty acid biomarkers allowed the identification of the main EPS producers of two contrasting types of intertidal marine sediments. Our study reveals that EPS production and degradation are supported by very different communities in muddy and sandy sediments and that EPS sources are more diverse in the sand. In mud, bound EPS are mainly derived from diatoms, while colloidal EPS are the result of degradation of bound exopolymers by certain specialised bacteria. In sand, bound EPS are rather of bacterial or cyanobacterial origin and diatoms contribute mainly to colloidal EPS. These differences are thought to be related to differences in the functioning of the epipelic and epipsammic communities and in particular to the use of EPS either for motility or for cell attachment purposes. 
\end {abstract}

\begin{keywords}
Extracellular Polymeric Substances | Stable isotopes | compound specific isotope analysis | fatty acids
\end{keywords}

$\ast$\begin{corrauthor}
cedric.hubas\at mnhn.fr
\end{corrauthor}

\section*{Introduction}
The term extracellular polymeric substances (EPS) is generic and refers to a wide variety of macromolecules whose main characteristic is to be of high molecular weight (> 10 kDa) and secreted by microbes outside the cell membrane. In intertidal sediments, these molecules are, for instance, secreted as a protection in response to changing environmental conditions or to allow cell motility \cite{tolhurst_diatom_2003}. But these secretions can also indirectly serve a number of ecosystem functions such as increasing the cohesion and adhesion properties of sediments \cite{lubarsky_stabilisation_2010}, or providing a significant source of organic matter at the base of the food web \cite{middelburg_fate_2000, bellinger_significance_2009}. They also represent a privileged pathway for cooperation between ecosystem engineers, leading to an improvement of the engineering effects on benthic communities \cite{passarelli_organisms_2014}.

Although many authors have studied these compounds and reviewed their multiple roles in aquatic ecosystems \cite{costerton_biofilms_1994, decho_microbial_2000,underwood_importance_2003,cescutti_bacterial_2010,wotton_essential_2004}, there is currently no clear classification.

As a general rule, exopolymers are classified into three categories which are basically distinguished by the proximity of the polymers to the membrane of the producing cells. This lack of homogeneity in the terms is most probably related to the high chemical diversity and complexity of EPS. 

Capsular polymer substances (CPS) are often defined as linked to the cell surface by a covalent bond to phospholipid or lipid A molecules, whereas EPS are released on the cell surface without being chemically attached to it and are often excreted to form a matrix more or less adherent to the surfaces \cite{cescutti_bacterial_2010}. EPS are further separated in two distinct fractions: bound-EPS which are tightly-bound long-chain material, and colloidal-EPS which are less refractory, small chain, easily extractable molecules. Colloidal EPS can be extracted by water at room temperature, while bound-EPS extraction requires hot water or bicarbonate \cite{underwood_importance_2003} or even cationic resins that trap the bivalent cations linking the EPS together, allowing the extraction of bound compounds \cite{takahashi_optimising_2009}. Thus, EPS are also sometimes described according to the extraction procedures. For instance, hot-bicarbonate and hot-water EPS (EPS$_{HB}$, EPS$_{HW}$), correspond to insoluble compounds solubilised using hot bicarbonate or water extraction protocols \cite{bellinger_biofilm_2005,hanlon_dynamics_2006}. 

These different EPS fractions differ in their biochemical composition \cite{passarelli_seasonal_2015} and it has been shown that different types of diatom-derived EPS drive changes in heterotrophic bacterial communities in intertidal sediments \cite{haynes_diatom-derived_2007,bohorquez_different_2017}.

The most significant progress on the subject concerns bacterial exopolysaccharides from microbial cultures (in particular pathogenic microorganisms), whose EPS metabolism and regulation mechanisms have been very well described. The genomic characterisation of these bacterial models of interest has led to fascinating discoveries. For example, it has been shown that EPS production (which underlies the development of bacterial biofilms) is under close control of a social behaviour called Quorum Sensing that allows interactions between members of microbial communities \cite{branda_biofilms_2005,solano_biofilm_2014}. Quorum sensing is based on the production and release of signalling molecules called autoinducers, which increase in concentration as a function of cell density \cite{camilli_bacterial_2006}. It was shown that these compounds were also present and particularly diverse in microbial mats \cite{decho_autoinducers_2009,decho_overview_2010}.

However, in the natural environment, the precise composition of EPS is still largely unknown. $^{13}$C-labelling experiment have highlighted the role of diatom organic matter as a growth substrate for benthic bacteria \cite{middelburg_fate_2000,veuger_fate_2012,oakes_composition_2010}. These studies traced diatom carbon and found that diatom EPS likely represents a link between benthic microalgae and higher trophic levels. Furthermore, the precise origin of these compounds in intertidal food webs is still subject to debate. Are diatoms the main, if not the only, producers of EPS in microphytobenthic assemblages, or do exopolymers present themselves rather as a pool of extracellular compounds of diverse origin?

In this study, we extracted colloidal and bound EPS from intertidal biofilms and analysed the natural stable isotope ratios (SIR) of carbon ($\delta^{13}$C) and nitrogen ($\delta^{15}$N). Isotope ratios of EPS were compared to those of fatty acid biomarkers to determine which microorganisms were primarily responsible for the production of EPS in muddy and sandy sediments.

Fatty acids are well recognised chemotaxonomic markers although they have a very limited taxonomic resolution and are hardly exclusive of a given organism \cite{de_carvalho_fatty_2014}. However, the ratios between the different fatty acids have shown convincing results in the identification and quantification of algal and bacterial groups and have already been successfully used to determine the composition of microbial mat \cite{dijkman_composition_2010} and sediment microbial communities \cite{findlay_efficacy_1989,passarelli_seasonal_2015,hubas_nanoscale_2017,gaubert-boussarie_untargeted_2020}.

The aim of this study was therefore to compare data of the natural stable isotopes of EPS with those of fatty acid biomarkers in two sediment types representative of intertidal environments (i.e. a muddy site and a sandy site), in order 1/ to accurately identify the main exopolymer producers and 2/ determine whether EPS production and dynamics was comparable between the microbial communities of contrasting sediment types.

\section*{Material and methods}

\subsection*{Sampling site}\label{m&m:site}

The sediment sampling took place in June 2017 at 2 tidal flat sites in France near La Coupelasse (Baie of Bourgneuf, France, Fig. \ref{figure1}). Bourgneuf Bay is a macrotidal bay located south of the Loire estuary  on  the  French  Atlantic  coast, containing large intertidal mudflats (100 \si{\km\squared}) colonized by microphytobenthic biofilms. The site is characterised by the extensive aquaculture of the Pacific oyster \textit{Crassostrea gigas}.  Oyster  farms  cover  about 10 \%  of  the  intertidal  area,  while  most  of  the  rocky  areas (about 17 \% of the intertidal area) are colonized by wild oysters \cite{le_bris_hyperspectral_2016} or macroalgae \cite{combe_mapping_2005,kazemipour_microphytobenthos_2012}.
Two contrasting sites were selected: a muddy site (47°0'53.326''N, 2°1'24.919''W) characterised by epipelic diatom communities and a high mud content (i.e. 50-90\%, \cite{meleder_spatio-temporal_2005}) and a sandy site (47°0'57.453''N, 2°1'33.676''W) characterised by epipsammic diatom communities and a low mud content (i.e. 40-60\%, \cite{meleder_spatiotemporal_2007}). The muddy site was sampled 4 times between $23^{th}$ and $28^{th}$ June 2017 (between 5 and 12 replicates depending on the date) while the sandy site was sampled 3 times (between 4 and 10 replicates)

\begin{figure*}
\centering
\includegraphics[scale=0.8]{figures/Fig1.pdf}
\caption{Studied area. (top panel) Map of Bourgneuf Bay (France) and location of the sampling sites.  left = true colors, right = Normalized difference vegetation index (NDVI); (bottom panel) sampling occasions according to the tidal level at the study sites (data provided by the Naval Hydrographic and Oceanographic Service (\href{https://maree.shom.fr/}{SHOM}) for coordinates: 047°06'00.0"N, 002°07' 00.0"W (Pornic). The first capital letter indicates the type of sediment (M=mud, S=sand), the other letters indicate the sampling point.}\label{figure1}
\end{figure*}

\subsection*{Exopolymeric substances (EPS)}\label{m&m:eps_ext}

Colloidal EPS were extracted by rotating sediment (for each sampling occasion, replicates of sediment core 10 \si{\centi\meter} diameter,  0.5 \si{\centi\meter} depth) in artificial sea water (salinity 30, Sea salts, NutriSelect® Basic) 1.5\si{\hour} at 4°C. Samples were then centrifuged (1500 g, 15 \si{\minute}) and the supernatant retrieved. Bound EPS were thereafter recovered by adding 2\si{\g} of a previously PBS-activated (4\si{\degree}C) Dowex Marathon C resin (sodium form, Sigma-Aldrich, Inc.) to the remaining pellet. After homogenisation, a second extraction was performed by rotating in artificial sea water 1.5 \si{\hour} at 4\si{\degree}C. Samples were then centrifuged (1500 g) again and the supernatant retrieved. Both supernatants form respectively the colloidal and bound fraction of EPS and were freeze-dried. Freeze-dried colloidal and bound EPS were weighted (in average $60 \pm 11$ \si{\milli\gram}) and the whole content was encapsulated in tin (Sn) capsules. They were placed in a 96 wells tray and analysed by  an Elementar Vario EL Cube or Micro Cube elemental analyzer (Elementar Analysensysteme GmbH, Hanau, Germany) interfaced to either an Isoprime VisION IRMS (Elementar UK Ltd, Cheadle, UK) or a PDZ Europa 20-20 isotope ratio mass spectrometer (Sercon Ltd., Cheshire, UK) by UC Davis Stable Isotope Facility. Samples were combusted at 1080\si{\degree}C in a reactor packed with chromium oxide and silvered copper oxide. Following combustion, oxides were removed in a reduction reactor (reduced copper at 650°C). The helium carrier then flows through a water trap (magnesium perchlorate and phosphorous pentoxide). CO$_2$ is retained on an adsorption trap until the N$_2$ peak is analyzed; the adsorption trap is then heated releasing the CO$_2$ to the IRMS.

In parallel, carbohydrate and protein concentrations were  measured following the phenol assay protocol \cite{dubois_colorimetric_1956} and the Lowry procedure \cite{lowry_protein_1951}, respectively. For carbohydrate analyses, 200 μl phenol (5\%) and 1 mL sulphuric acid (98\%) were added to 200 μl of previously extracted colloidal and bound supernatants. They were then incubated for 35 min at 30\si{\degree}C and the carbohydrate concentration was measured using a spectrophotometer (Milton Roy Spectronic Genesys 2). The optical density of the solution was measured at 488 nm. For protein analyses, 250 μl subsamples were incubated for 15 min at 30\si{\degree}C with 250 μl of 2\% sodium dodecyl sulphate salt (SDS) and 700 μl of a chemical reagent prepared as described in \cite{lowry_protein_1951}. The subsamples were then incubated for another 45 min at 30\si{\degree}C with 100 μl of Folin reagent (diluted with distilled water 5:6 v/v). The protein concentration was measured by spectrophotometry at 750 nm. Calibration curves were prepared using glucose and bovine serum albumin (BSA) as standards for carbohydrates and proteins, respectively.

\subsection*{Fatty acid extraction}\label{m&m:FA_ext}

Fatty acid (FA) analysis was performed on triplicates of sediment core 10 \si{\centi\meter} diameter,  0.5 \si{\centi\meter} depth) following the method of \cite{bligh_rapid_1959} as modified by \cite{meziane_fatty_2000} and \cite{passarelli_seasonal_2015}. Lipids were extracted with a 20 \si{\minute} ultrasonication (sonication bath, 80 \si{\kilo\hertz}, Fisherbrand™) in a mixture of distilled water, chloroform and methanol in ratio 1:1:2 (v:v:v, in \si{\milli\liter}). Lipids were concentrated under $N_2$ flux, and saponified, in order to separate FA, with a mixture of NaOH (2 \si{\mole\per\liter}) and methanol (1:2, v:v, in \si{\milli\liter}) at 90 \si{\degree}C during 90 \si{\minute}. Saponification was stopped with 500 \si{\micro\liter} hydrochloric acid. Samples were then incubated with BF3-methanol at 90 \si{\degree}C during 10 \si{\minute} to transform free fatty acids into fatty acid methyl esters (FAME), which were isolated and kept frozen in chloroform. Just before analysis, samples were dried under $N_2$ flux and transferred to hexane.

\subsection*{Compound specific isotope analysis (CSIA) of FAME}\label{m&m:CSIA}

Carbon stable isotope ratios (expressed in \textperthousand) of individual fatty acids were measured by gas-chromatography-isotope ratio mass spectrometry (GC-IRMS). Measurements were performed at the Stable Isotope Platform of the European Institute for Marine Studies (IUEM, Brest, France). FAMEs were injected in splitless mode and separated using a B5HT column (30  m × 0.25  mm ID × 0.2 μm, Phenomenex) with a Thermo Fisher Scientific TRACE GC ULTRA equipped with GC isolink combustion, Conflo IY interace and Delta V plus (Thermo Fisher Scientific) isotope ratio mass spectrometer (IRMS). Fatty acids were converted into CO$_2$ by combustion in the ISOLINK furnace and transferred to the CONFLO IV interface and then introduced to the IRMS. Fatty acid methyl esters were identified by comparison of their retention time with those of commercial standards and in-house standard mixtures. Both FA 18:1n-9 and 18:3n-3 co-eluted and were analysed simultaneously. Fatty acids kept for $\delta^{13}$C analyses were selected based on their abundance and detection in CSIA (i.e., with amplitudes > 800 mV). Stable carbon isotope ratios for individual FA were calculated from FAME data by correcting for the one carbon atom in the methyl group that was added during the derivatization process. This correction was made according to \cite{gladyshev_stable_2012} by taking into account the isotope ratio of the derivatized methanol (BF3 methanol), and the fractional carbon contribution of the free FA to the ester.

\begin{equation}
\label{eq1}
    \delta^{13}\text{C}_{FA}=\frac{(\delta^{13}\text{C}_{FAME}-(1-f)\delta^{13}\text{C}_{CH_3OH})}{f}
\end{equation}

where $\delta^{13}\text{C}_{FA}$ and $\delta^{13}\text{C}_{FAME}$ (in \textperthousand) are the isotopic composition of the free FA, and the FA methyl ester respectively, $f$ is the fractional carbon contribution of the free FA to the ester and $\delta^{13}\text{C}_{CH_3OH}$ is the isotopic composition of the methanol derivatization reagent (–39.1 \textperthousand).

\subsection*{Fatty acid identification}\label{m&m:FA_id}

Identification of the samples was performed using a gas chromatograph coupled to mass spectrometer (GC–MS, Varian 450GC with Varian 220-MS). Compounds annotation was performed by comparing mass spectra with NIST 2017 library. Corresponding fatty acids are designated as X:Yn-Z, where X is the number of carbons, Y the number of double bonds and Z the position of the ultimate double bond from the terminal methyl.

\subsection*{Statistical analyses}\label{m&m:stats}

Univariate statistics were carried out by checking the normality of the data per group (Shapiro test) and the homogeneity of the variances (Bartlett or Levene test). Where the data did not meet these criteria or the sample size were too small, we applied van der Waerden normal scores test followed by Fisher's least significant difference (LSD) post-hoc test.
In case the sample size was larger but the conditions were still not met, we used Permutation one-way Welch's Anova followed by Tukey HSD posthoc test. In case we had to compare two samples, we checked for normality and equality of variance (Fisher-Snedecor test) and used Welch’s permutation t-test or student t-test. All analyses were performed using R version 4.0.3 using the "stats" package.
We performed a smoothed density estimation on the fatty acid isotope ratio data using the geom\textunderscore smooth function of the "ggplot2" package. The function, computed and drawn kernel density estimate based on the observed distribution of the stable isotopes ratio.
Data are available at \href{https://doi.org/10.5281/zenodo.7351530}{https://doi.org/10.5281/zenodo.7351530}. Statistical scripts and command lines are available on \href{https://github.com/Hubas-prog/EPS_FA_CSIA}{GitHub} at the following address \href{https://doi.org/10.5281/zenodo.7387066}{https://doi.org/10.5281/zenodo.7387066}.

\section*{Results and discussion}

In the present study, compound-specific isotope analysis (CSIA) of fatty acid biomarkers was used to infer about possible origin of microbial EPS. Our main assumption was that isotopic fractionation between the microoganisms and the product of their metabolism (i.e. EPS, fatty acids) is null or negligible. At present, no study has been able to demonstrate with certainty whether this hypothesis is true or false. There is, however, evidence that fractionation exists between microorganisms and their food sources. In bacteria, substantial isotopic fractionation has been shown between biomarker lipids and their growth substrate \cite{summons_carbon_1994} with bacterial biomarkers being significantly depleted in $ ^{13}$C compared to the food source. In \textit{Escherichia coli}, respired CO$_2$ was 3.4\textperthousand$ $ depleted in $ ^{13}$C relative to glucose (used as the carbon source) although total cellular carbon was only 0.6\textperthousand$ $ depleted in $ ^{13}$C, and lipid fractions by 2.7\textperthousand$ $ \cite{blair_carbon_1985}. But to date however, there is no evidence in the literature that the same phenomenon exists between microorganisms and their metabolites. 

\subsection*{Elemental EPS compositions}\label{Res:EPS_compo}

Carbon and nitrogen contents were significantly different between sites as well as between bound and colloidal EPS (table \ref{tab:table1}, Fig. \ref{figure2}). Bound EPS were almost always richer in carbon and nitrogen than colloidal EPS (fig. \ref{figure2}a,c). The only noticeable exception was at Ms1. We also noted a very significant decrease in the N and C contents in the colloidal fraction at this site (i.e. muddy site) between Ms1 and Ms2. Both were sampled the same day but respectively at ebbing and rising tide. These findings are partly consistent with those of Hanlon et al. \cite{hanlon_dynamics_2006}. During periods of diurnal emersion at a muddy site, these authors reported that bacteria converted bound EPS into more labile colloidal EPS. By analogy, we can hypothesise that bacteria at our site were very efficient at converting bound EPS to colloidal EPS (hence the slight decrease in N and C content in bound EPS) but that they probably also consume colloidal EPS at very high rates.

The same patterns were observed for sugar and protein concentrations measured by colorimetry, but with greater variability between measurements (supplementary figure \ref{figureS1}). If we focus on colloidal EPS, we notice that the consumption of these between Ms1 and Ms2 mainly concerned carbohydrates. Exopolymers are mainly composed of carbohydrates and proteins \cite{wotton_essential_2004} which therefore represent the main sources of C and N in EPS. Overall, bacterial EPS contain more proteins and higher molecular diversity than diatomaceous EPS \cite{decho_bacterial_1990}. The carbohydrates produced by microphytobenthos are mainly heteropolymers, with a large diversity of molecules. They range in molecular weight from few monosaccharides to highly complex molecules whose relative proportion in terms of monomers determines the physicochemical structure and hydrophobic characteristics of the EPS matrix \cite{underwood_importance_2003,passarelli_seasonal_2015}. The higher C content of the EPS is therefore probably partly related to a higher proportion of sugars of diatom origin.

\begin{table}[!t]
\caption{Comparison of Carbon and Nitrogen contents and isotopic ratios of colloidal and bound EPS in mud and sand using the van der Waerden (Normal Scores) non parametric test. df = degree of freedom. Results of the post-hoc test using the criterium Fisher's least significant difference (LSD) are shown in Fig. \ref{figure2}}
\label{tab:table1}
\begin{tabular}{lllll}
\hline
Variable & $\chi^2$ & df & p-value   \\ \hline
Carbon content  & 46.83024   & 13  & 1.03209e-05 \\
Nitrogen content    & 27.364   & 14  & 6.74533e-06 \\
    $\delta^{13}$C  & 36.31005   & 13  & 0.00053  \\
$\delta^{15}$N  & 35.02141   & 13  & 0.00084  \\ \hline
\end{tabular}%
\end{table}

\subsection*{EPS isotopic compositions}\label{Res:EPS_iso}

At the muddy site, bound EPS were always $^{13}$C or $^{15}$N depleted in comparison to colloidal EPS at this site (Fig. \ref{figure2}b,d). At the sandy site, the same pattern is observed but both nitrogen and carbon stable isotope ratio showed a higher variability.

All sampling dates together, $\delta^{13}$C (Fig. \ref{figure3}a, top panel) and $\delta^{15}$N values  were significantly different between bound and colloidal EPS at the muddy site (Permutation two Sample t-tests, $\delta^{13}$C: t = -10.678, p-value = 0.002; $\delta^{15}$N: t = -4.4325, p-value = 0.002).

At the sandy site, $\delta^{13}$C values were also significantly different (Fig. \ref{figure3}a, bottom panel) between bound and colloidal EPS (two Sample Student's t-test, t = -4.9474, df = 22, p-value = 5.984e-05) but $\delta^{15}$N was not significantly different (two Sample Student's t-test, t = -0.97547, df = 22, p-value = 0.3399).

\begin{figure*}[t]
\centering
\includegraphics[scale=0.6]{figures/Fig2.pdf}
\caption{Chemical composition of the EPS. \textbf{a},\textbf{c}: Carbon (C) and Nitrogen (N) contents in \si{\micro\gram} per \si{\milli\gram} of freeze dried EPS. \textbf{b},\textbf{d}: Carbon and Nitrogen stable isotope ratio ($\delta$ notation against atmospheric $N_2$ and Vienna PDB respectively) of the EPS. Colloidal EPS corresponded to loose, water-extractable exopolymers whereas bound EPS correspond to ion exchange resin-extractable exopolymers. Letters within the graph represent results of Fisher’s least significant difference (LSD) post-hoc test. For the corresponding van der Waerden test, please see Table \ref{tab:table1}}\label{figure2}
\end{figure*}

All sampling dates grouped together $\delta^{13}$C between bound and colloidal EPS were thus always significantly different at both sites (Fig. \ref{figure3}a), indicating that these two fractions were from different EPS producers. Comparison with the literature is difficult as it is the first time that C and N natural stable isotopes ratio are reported on intertidal bound and colloidal EPS. The values reported in the literature for the main monosaccharides constituting the extracellular sugars are however in agreement with our results (i.e. a natural $\delta^{13}$C of -15 to -18 $\text{\textperthousand}$) \cite{oakes_composition_2010}

\subsection*{Carbon isotope ratio of fatty acid classes}\label{res:FA_class_compo}

In sandy sediment $\delta^{13}$C were significantly different between fatty acids classes ($F=23.128, df1=3, df2=109, p= \num{1.16e-11}$) and showed a gradual $^{13}$C enrichment (Fig. \ref{figure3}b) from branched fatty acids (BFA) to mono- (MUFA) and poly-unsaturated (PUFA) fatty acids. Such differences were not observed in the muddy site. In the mud, $\delta^{13}$C of BFA, saturated (SFA) and MUFA were not significantly different. Only PUFA showed a slightly higher mean $\delta^{13}$C (Permutation one-way Welch Anova followed by Tukey HSD posthoc test, $F=33.588, p < \num{2.2e-16}$).

\begin{figure}
\centering
\includegraphics[scale=0.35]{figures/Fig3.pdf}
\caption{Comparison of $\delta^{13}$C (corrected according to equation \ref{eq1}) between fatty acid classes and EPS fractions. \textbf{a}: bound and colloidal EPS were significantly different (t-tests, p < 0.001) at both sites. \textbf{b}: numbers indicate significantly different groups as evidenced by post-hoc tests. BFA=branched, SFA=saturated, MUFA=monounsaturated and PUFA=polyunsaturated fatty acids}\label{figure3}
\end{figure}

In comparison with similar ecosystems (i.e. intertidal muddy sediments), the isotope ratios of the main fatty acids are quite consistent. Previous studies recorded $\delta^{13}$C ranging from -16 to -21$\text{\textperthousand}$ for branched, -14 to -26$\text{\textperthousand}$ for saturated, -13 to -22$\text{\textperthousand}$ for monounsaturated and -15 to -22$\text{\textperthousand}$ for polyunsaturated fatty acids \cite{taylor_microphytobenthic_2013,veuger_fate_2012,middelburg_fate_2000}. Taylor et al. \cite{taylor_microphytobenthic_2013} also showed that natural carbon isotope ratios were highly variable even over relatively short periods (i.e. 30h). These changes indicate that subtle modifications in the metabolic processes of carbon assimilation as well as interactions between microorganisms can take place over very short periods and could explain the variability of our $\delta^{13}$C values.

The tetracosanoic acid (SFA, 24:0) was excluded from the above mentioned analyses as it increased dramatically the variability because of extreme and unusually negative $\delta^{13}$C values indicative of a specific metabolism. The mean $\delta^{13}$C of 24:0 was $-66.89 \pm 35.84 \text{\textperthousand}$ and $-59.24 \pm 71.82 \text{\textperthousand}$ in the mud and sand respectively. It also sometimes showed a plurimodal distribution (as shown by density plots figure \ref{figure4}b) which indicate that 24:0 had likely varied microbial origins. This particular fatty acid was the only one to show extremely low $\delta^{13}$C values in line with the isotopic ratios generally found in methane-rich ecosystems for which direct links could be established between $\delta^{13}$C values and the presence of methane-oxidizers in bacterial communities 
\cite{paul_methane-oxidizing_2017,uhlig_methane-oxidizing_2018}. It is indeed possible that the 24:0 originated from anaerobic bacteria related to the oxidation of methane or the sulphur cycle. The most negative $\delta^{13}$C values were recorded in highly reduced muddy sediments. Unfortunately, it is not possible to establish a direct link in our study.

\begin{figure*}
\centering
\includegraphics[scale=0.65]{figures/Fig4.pdf}
\caption{C and N isotopic ratio of fatty acids and EPS fractions. \textbf{a-d}: Kernel density estimates of $\delta^{13}$C of fatty acid biomarkers (corrected according to equation \ref{eq1}). \textbf{e}:  $\delta^{13}$C and  $\delta^{15}$N biplots of bound and colloidal EPS. All sampling points were grouped together. In panels \textbf{a-d}, fatty acids are grouped by classes they belong to according to fig.\ref{figure3}}\label{figure4}
\end{figure*}

\subsection*{Biomarkers revealed contrasting EPS producers between sites}\label{res:_FA_bio_compo}

The $\delta^{13}$C values of fatty acid biomarkers were measured at each site. By comparing the distribution of individual fatty acids (Fig. \ref{figure4}a-d) with the carbon isotope ratio of the EPS (Fig. \ref{figure4}e), it was relatively straightforward to determine which fatty acids had an isotope ratio closest to that of bound and colloidal EPS. Based on figure \ref{figure4} and taking into account the quality of the alignment between fatty acids and EPS, a detailed literature review of the potential origins of EPS in the studied sediments was performed. The result is available in the supplementary table \ref{tab:table2}. The analysis revealed that EPS producers were very different between the two sites.
In the mud, colloidal EPS were potentially mainly produced by bacteria, whereas bound EPS were mainly produced by diatoms with a significant contribution from cyanobacteria and bacteria.
In the sand, the origins of EPS were more diversified. Colloidal EPS were mainly produced by diatoms and bacteria with a potential contribution from cyanobacteria. Bound EPS were mainly produced by bacteria. The sediments of the study site are, indeed, known to harbour microphytobenthic assemblages dominated by diatoms (i.e. 97\% \cite{meleder_spatiotemporal_2007}). Depending on the site, these are accompanied by cyanophyceae, euglenophyceae and chlorophyceae \cite{meleder_spatiotemporal_2007}.

\subsection*{Epipelic and epipsammic diatoms contributed differently to the EPS chemistry}\label{res:_diatom-contrib}

Most common fatty acids in diatoms are myristic acid (14:0), palmitic acid (16:0), palmitoleic acid (16:1n-7), docosahexaenoic acid (DHA, 22:6n-3), and eicosapentaenoic acid (EPA, 20:5n-3) \cite{yi_exploring_2017}. In terms of relative proportions, however, 16:1n-7 and 20:5n-3 generally dominate the total fatty acids \cite{dunstan_essential_1994}. These two fatty acids had relatively close $\delta^{13}$C values that best aligned respectively with bound EPS in muddy sediments ($-20.3 \pm 1.1 \text{\textperthousand}$) and with colloidal EPS ($-13.4 \pm 4.5 \text{\textperthousand}$) in sandy sediments. This indicated a very different functioning between the assemblages at these two sites.

The muddy site had an epipelic community typical of these environments (presence of characteristic migratory behaviour, pers. obs.) whereas the sandy site has all the characteristics of epipsammic communities \cite{meleder_spatiotemporal_2007}. Thus it appears that epipelic diatoms mainly contributed to the bound EPS fraction while epipsammic diatoms mainly contributed to the colloidal EPS pool. This differential contribution according to habitat can be explained by the implementation of different adaptation strategies of diatoms to environmental parameters.

Epipelic diatoms secrete large quantities of extracellular exopolymers that are involved in motility. Mucilage is secreted from the raphe and adheres to the sediment following hydration. Cellular movement is then generated when the EPS associated with the trans-membrane complexes is displaced along the raphe line by actine microfilament bundles \cite{underwood_importance_2003,cartaxana_effects_2011}. The products necessary for the migration of the diatoms are therefore secreted and used in the immediate vicinity of the cell. This is most probably the reason why we observed a massive contribution of diatoms to bound EPS at the muddy site.

In a previous study, our team measured the monosaccharide compositions of sandy intertidal sediment EPS \cite{passarelli_seasonal_2015}. As a result of the accumulation of silt in these sediments (caused by the implantation of biogenic structures), and the evolution of the diatom assemblage towards an epipelic community, we observed a modification of the sugars produced, which only occurred in the bound fraction. This further confirms the large contribution of epipelic diatoms to the bound EPS pool of muddy sediments.

In contrast, epipsammic diatoms mainly contributed to colloidal extracellular polymers together with cyanobacteria, green algae and bacteria. Epipsammic diatoms do not migrate because they live in sediments which are very dynamic and which have a low light extinction coefficient over achievable distances of the order of hundreds of micrometers \cite{cartaxana_physiological_2011}. As a result, these diatoms do not migrate but instead used adhesion to sand particle to avoid being resuspended. In the absence of photomigratory response, they much more rely on strong photophysiological protection mechanisms than epipelic motile diatoms \cite{cartaxana_physiological_2011}. Capsular and bound EPS were thus instead rather produced sparingly and used for attachment and fixation purposes.

In a benthic freshwater diatom, it has been shown that capsular EPS mainly consist of glycoprotein that develop from fibrillar precursors and that bacteria preferentially attach to encapsulated diatom cells \cite{leinweber_capsules_2015}. This is probably a strategy of the bacteria to maximise the chances of success in terms of adhesion and also to ensure access to an important food source. This may explain why bound EPS were mainly aligned with bacterial biomarkers at the this site.

\subsection*{Multiple EPS origins favour the development of EPS-specialised bacteria}\label{res:_FA_decompo}

Since bound EPS best aligned with branched fatty acids, 18:1n-7 and some SFA at the sandy site (Table \ref{tab:table2}), we could conclude that bound EPS were mainly of bacterial origin \cite{kharlamenko_fatty_1995,middelburg_fate_2000,perry_fatty_1979,hubas_proliferation_2013,taylor_microphytobenthic_2013,boon_significance_1977} at this site either as a direct production or as a result of degradation of bound and capsular diatomic EPS. Therefore, diatoms mainly contributed directly to the colloidal fraction which was also degraded by specialised bacteria (as shown by 18:1n-7).

It is very difficult and even impossible to assign a given branched fatty acid to a specific bacterial taxon. Certain fatty acids may represent a significant proportion of total fatty acids in certain bacterial groups or taxa. Vaccenic acid (18:1n-7), for example, can account for more than 30\% of the total in purple bacteria \cite{kharlamenko_fatty_1995}. Similarly, 15:0iso and 15:0anteiso fatty acids may be dominant in \textit{Desulfovibrio sp.} species \cite{boon_significance_1977}. But only a limited number of
bacteria have unusual fatty acids. By contrast, branched-chain fatty acids of the iso and anteiso series occur widely in bacteria, give a complex pattern, and are therefore valuable in bacterial systematics \cite{kaneda_iso-_1991}. In the present study, it is therefore the changes in the relative composition and/or dominance of bacterial fatty acids within the different EPS fractions that indicated changes in microbial assemblages, rather than the presence of any particular fatty acid.

In addition, we also observed that branched fatty acids 15:0anteiso and 15:0iso (Fig. \ref{figure4}a) showed a bimodal distribution of their $\delta^{13}$C value at the sandy site. This can be explained by the fact that these fatty acids originated from different bacterial species with different C sources (i.e. bound vs. colloidal EPS) and further confirm the existence of prokaryotic assemblages dedicated to each EPS fractions.

Earlier $^{13}$C enrichment experiments have already shown EPS consumption by bacteria through 15:0anteiso and 15:0iso enrichment but also provided additional evidence that some taxa (e.g. Acinetobacter) might be considered specialist EPS-degrading bacteria \cite{taylor_microphytobenthic_2013}.

Similarly, the presence of "EPS degraders" can also be demonstrated at the muddy site. At this site, colloidal EPS aligned well with 17:0iso indicating that specific taxa rich in this branched fatty acid are predominantly involved in the production of colloidal EPS, probably from the degradation of diatom bound EPS.

\section*{Conclusions}
By comparing the natural C and N stable isotope ratios of fatty acids and bound and colloidal EPS fractions in intertidal sediments, we identified a very different dynamics of EPS production and degradation between sandy and muddy sites. The most noticeable difference was that epipelic and episammic diatoms contributed differently to the chemistry of the EPS, which had an important implication for the development of EPS specialised bacteria. These differences are thought to be related to differences in the functioning of the epipelic and epipsammic communities and in particular to the use of EPS either for motility or for cell attachment purposes. 

\begin{Conflict}
The authors declare that they have no financial conflicts of interest in relation to the content of the article.
\end{Conflict}

\begin{contributions}
Conceptualization: KS, BJ, CH ; Data Curation: CH ; Formal analysis: CH ; Funding acquisition: KS, BJ, CH ; Investigation: all authors ; Methodology: all authors ; Project administration: KS, BJ, CH ; Resources: all authors ; Software: CH ; Supervision: KS ; Validation: all authors ; Visualization: CH ; Writing – original draft: CH ; Writing – review \& editing: CH, BJ, VM, KS.
\end{contributions}

\begin{acknowledgements}
Some data were acquired on research platforms, which we would like to thank. CSIA data were acquired at Stable Isotope Platform of the European Institute for Marine Studies (IUEM, Brest, France). EA-IRMS data were acquired at the UC Davis stable isotope facility.
\end{acknowledgements}

\begin{funding}
This study was supported by the BIO-Tide project (The role of microbial biodiversity in the functioning of marine tidal flat sediments), funded through the 2015–2016 BiodivERsA COFUND call for research proposals, with the national funders BelSPO (BRAIN-be contract BR/175/A1/BIO-Tide-BE), FWO (3G0H6816), ANR (ANR-16-EBI3-0008), and SNSF. The post-doctoral grant of J. G-B was supported by the Regional Council of Brittany, SAD program and META-Tide projects.
\end{funding}

\section*{Bibliography}
\bibliography{EPSarticle.bib}

%% You can use these special %TC: tags to ignore certain parts of the text.
%TC:ignore
%the command above ignores this section for word count

% lire : https://aslopubs.onlinelibrary.wiley.com/doi/epdf/10.4319/lo.2013.58.4.1463
% Lire : https://www.sciencedirect.com/science/article/pii/S1385110113001962?casa_token=Am6ZMKQQuMQAAAAA:r3GGxE-Cg2ZEHfg-p4N4_MiovmYNTEcuICPH7NsQ1FCWduWRMkTOxV7o46t1Fk956oaQQVwwDA
%lire : https://aslopubs.onlinelibrary.wiley.com/doi/pdfdirect/10.4319/lo.2010.55.5.2126


\onecolumn
\newpage


%%%%%%%%%%%%%%%%%%%%%%%%%%%%%
% Supplementary Information %
%%%%%%%%%%%%%%%%%%%%%%%%%%%%%
\beginsupplement
%\captionsetup*{format=largeformat}
%\section{Supplementary information} \label{SF1} 

%SUPP FIG
\begin{figure}[]
\centering
\includegraphics[scale=0.5]{figures/FigSF1.pdf}
\caption{Colorimetric measurements of EPS concentrations in \si{\milli\gram} equivalent to Glucose or Bovine Serum Albumine (BSA) for carbohydrates and proteins respectively per \si{\milli\liter} of extracted EPS; colloidal EPS corresponded to loose, water-extractable exopolymers whereas bound EPS correspond to ion exchange resin-extractable exopolymers. }\label{figureS1}
\end{figure}

% SUPP TABLE
\begin{table}[]
\begin{centering}
\caption{Presumed sources of colloidal and bound EPS (carbohydrates, proteins) at muddy and sandy sites. Position refers to the quality of alignment between fatty acids and EPS $\delta^{13}$C values: Aligned = the mean $\delta^{13}$C value of a given fatty acid was within the standard deviation or confidence interval of the corresponding EPS isotope ratio, Sup. (Superimposed)= $\delta^{13}$C values of EPS and fatty acids overlapped by their standard deviations or confidence intervals. \underline{\textbf{Bold underlined}} = major fatty acid (20-40\%) in the corresponding sources. \textbf{Bold}  = important fatty acid (10-20\%), \textit{Italic} = present in trace amounts (<10\%)}
\label{tab:table2}
\begin{tabular}{@{}lllll@{}}
\toprule
Location & EPS type & Fatty acids & Position & Possible origin of FA \\ \midrule
\multicolumn{1}{c}{\multirow{15}{*}{Mud}} & \multirow{4}{*}{Colloidal} & 22:6n-3 & \multirow{2}{*}{Aligned} & \underline{\textbf{Dinoflagellates}}, \textbf{Haptophyta} \cite{jonasdottir_fatty_2019}, \textit{Diatoms, Cyanobacteria} \cite{jonasdottir_fatty_2019,dunstan_essential_1994,dijkman_composition_2010}  \\
\multicolumn{1}{c}{} &  & 20:0 &  & \textit{Bacteria} \cite{pegu_isolation_2020,perry_fatty_1979} \\ \cmidrule(l){3-5} 
\multicolumn{1}{c}{} &  & 18:4n-3 & \multirow{2}{*}{Sup.} & \textbf{Haptophyta}, \textbf{Pheophyceae} \cite{guerrero_microbial_2002}, \textit{Diatoms} \cite{dunstan_essential_1994} \\
\multicolumn{1}{c}{} &  & 17:0iso &  & \underline{\textbf{Bacteria}} \cite{perry_fatty_1979,kaneda_iso-_1991} \\ \cmidrule(l){2-5} 
\multicolumn{1}{c}{} & \multirow{11}{*}{Bound} & 20:4n-6 & \multirow{3}{*}{Aligned} & \textit{Diatoms} \cite{dunstan_essential_1994,dijkman_composition_2010,jonasdottir_fatty_2019}, \textit{Chlorophyta} \cite{Jonasdottir2019FattyPhytoplankton} \\
\multicolumn{1}{c}{} &  & 17:1n-5/7 &  & \textit{Bacteria} \cite{pegu_isolation_2020} \\
\multicolumn{1}{c}{} &  & 16:4n-1 &  & \textbf{Diatoms} \cite{leu_ultraviolet_2007}, \textit{Diatoms} \cite{jonasdottir_fatty_2019,dunstan_essential_1994} \\ \cmidrule(l){3-5} 
\multicolumn{1}{c}{} &  & 22:0 & \multirow{8}{*}{Sup.} & \textit{Diatoms} \cite{leu_ultraviolet_2007}, \textit{Cyanobacteria} \cite{sahu_fatty_2013} \\
\multicolumn{1}{c}{} &  & 20:5n-3 &  & \underline{\textbf{Diatoms}} \cite{leu_ultraviolet_2007,dunstan_essential_1994,dijkman_composition_2010}, \textbf{Diatoms} \cite{jonasdottir_fatty_2019} \\
\multicolumn{1}{c}{} &  & 18:4n-6 &  & \textit{Cyanobacteria} \cite{vyssotski_chemical_2015} \\
\multicolumn{1}{c}{} &  & 18:2n-6 &  & \textbf{Cyanobacteria, Chlorophyta} \cite{sahu_fatty_2013,jonasdottir_fatty_2019}, \textbf{Fungi} \cite{vallet_laminariales_2020}  \\
\multicolumn{1}{c}{} &  & 18:1n-9/18:3n-3 &  & \underline{\textbf{Cyanobacteria, Chlorophyta}}  \cite{sahu_fatty_2013,jonasdottir_fatty_2019} /  \cite{dijkman_composition_2010,jonasdottir_fatty_2019} * \\
\multicolumn{1}{c}{} &  & 17:0 &  & \textit{Bacteria, Diatoms} \cite{perry_fatty_1979,leu_ultraviolet_2007}\\
\multicolumn{1}{c}{} &  & 17:iso &  & \underline{\textbf{Bacteria}} \cite{perry_fatty_1979,kaneda_iso-_1991} \\
\multicolumn{1}{c}{} &  & 16:3n-4/16:4n-3 &  & \textit{Diatoms} \cite{jonasdottir_fatty_2019,remize_study_2020}/ \textbf{Chlorophyta} \cite{jonasdottir_fatty_2019} \\
\multicolumn{1}{c}{} &  & 15:0 &  & \underline{\textbf{Bacteria}} \cite{pegu_isolation_2020}, \textit{Diatoms, Chlorophyta} \cite{dijkman_composition_2010,sahu_fatty_2013,dunstan_essential_1994} \\ \midrule
\multirow{21}{*}{Sand} & \multirow{15}{*}{Colloidal} & 22:6n-3 & \multirow{13}{*}{Aligned} & \underline{\textbf{Dinoflagellates}}, \textbf{Haptophyta} \cite{jonasdottir_fatty_2019}, \textit{Diatoms, Cyanobacteria} \cite{jonasdottir_fatty_2019,dunstan_essential_1994,dijkman_composition_2010} \\
 &  & 20:5n-3 &  & \underline{\textbf{Diatoms}} \cite{leu_ultraviolet_2007} \\
 &  & 20:4n-6 &  & \textit{Diatoms} \cite{dunstan_essential_1994,dijkman_composition_2010,jonasdottir_fatty_2019}, \textit{Chlorophyta} \cite{jonasdottir_fatty_2019} \\
 &  & 18:4n-6 &  & \textit{Cyanobacteria} \cite{vyssotski_chemical_2015} \\
  &  & 18:2n-6 &  & \textbf{Cyanobacteria, Chlorophyta} \cite{sahu_fatty_2013,jonasdottir_fatty_2019} \\
 &  & 18:1n-9/18:3n-3 &  & \underline{\textbf{Cyanobacteria, Chlorophyta}}  \cite{sahu_fatty_2013,jonasdottir_fatty_2019} /  \cite{dijkman_composition_2010,jonasdottir_fatty_2019} * \\
 &  & 18:1n-7 &  & \textbf{Bacteria} \cite{middelburg_fate_2000} \\
 &  & 17:1n-5/7 &  & \textit{Bacteria} \cite{pegu_isolation_2020}  \\
 &  & 17:0 &  & \textit{Bacteria, Diatoms} \cite{perry_fatty_1979,leu_ultraviolet_2007} \\
 &  & 16:3n-4/16:4n-3 &  & \textit{Diatoms} \cite{jonasdottir_fatty_2019,remize_study_2020}/ \textbf{Chlorophyta} \cite{jonasdottir_fatty_2019} \\
 &  & 16:1n-7 &  & \underline{\textbf{Diatoms}} \cite{leu_ultraviolet_2007,dunstan_essential_1994}, \textbf{Cyanobacteria, Bacteria} \cite{jonasdottir_fatty_2019,dijkman_composition_2010,sahu_fatty_2013,perry_fatty_1979}, \textit{Chlorophyta} \cite{sahu_fatty_2013} \\
 &  & 16:1n-5 &  & \underline{\textbf{Diatoms, Bacteria}} \cite{perry_fatty_1979,leu_ultraviolet_2007,dunstan_essential_1994} \\
 &  & 16:0 &  & Major or important fatty acid in various sources \cite{perry_fatty_1979,leu_ultraviolet_2007,dunstan_essential_1994,dijkman_composition_2010,sahu_fatty_2013,guerrero_microbial_2002,jonasdottir_fatty_2019,vallet_chemically-mediated_2018} \\
 &  & 15:0 &  & \underline{\textbf{Bacteria}} \cite{pegu_isolation_2020}, \textit{Diatoms, Chlorophyta} \\ \cmidrule(l){3-5} 
 &  & 14:0 & \multirow{2}{*}{Sup.} & \textbf{Diatoms} \cite{leu_ultraviolet_2007} \\
 &  & 18:0 &  & \textbf{Dinoflagellates} \cite{jonasdottir_fatty_2019} * \\ \cmidrule(l){2-5} 
 & \multirow{6}{*}{Bound} & 15:0iso, 15:0anteiso & Aligned & \underline{\textbf{Bacteria}} \cite{boon_significance_1977}, \textit{Bacteria} \cite{perry_fatty_1979,taylor_microphytobenthic_2013,middelburg_fate_2000} \\ \cmidrule(l){3-5}
 &  & 18:1n-7 & \multirow{5}{*}{Sup.} & \textbf{Bacteria} \cite{kharlamenko_fatty_1995,middelburg_fate_2000,perry_fatty_1979,hubas_proliferation_2013,taylor_microphytobenthic_2013}, \textit{Cyanobacteria, Chlorophyta, Diatoms} \cite{dunstan_essential_1994,jonasdottir_fatty_2019,dijkman_composition_2010} \\
 &  & 18:0 &  & \textbf{Dinoflagellates} \cite{jonasdottir_fatty_2019} * \\
 &  & 16:0 &  & Major or important fatty acid in various sources \cite{perry_fatty_1979,leu_ultraviolet_2007,dunstan_essential_1994,dijkman_composition_2010,sahu_fatty_2013,guerrero_microbial_2002,jonasdottir_fatty_2019,vallet_chemically-mediated_2018} \\
 &  & 15:0 &  & \underline{\textbf{Bacteria}} \cite{pegu_isolation_2020}, \textit{Diatoms, Chlorophyta} \\
 &  & 14:0 &  & \textbf{Diatoms} \cite{leu_ultraviolet_2007} \\ \bottomrule
 \end{tabular}
* also detected in all sources in trace amounts \cite{vallet_laminariales_2020,leu_ultraviolet_2007,jonasdottir_fatty_2019,perry_fatty_1979,guerrero_microbial_2002,dunstan_essential_1994,pegu_isolation_2020,dijkman_composition_2010,sahu_fatty_2013}.
\end{centering}
\end{table}


%TC:endignore
%the command above ignores this section for word count

\end{document}
