## Supplementary material for "Identification of microbial exopolymer producers in sandy and muddy intertidal sediments by compound-specific isotope analysis": Latex_zip_file_V2: EPSarticle.bib

@article{VANHEUKELEM200131,
title = {Computer-assisted high-performance liquid chromatography method development with applications to the isolation and analysis of phytoplankton pigments},
journal = {Journal of Chromatography A},
volume = {910},
number = {1},
pages = {31-49},
year = {2001},
issn = {0021-9673},
doi = {https://doi.org/10.1016/S0378-4347(00)00603-4},
url = {https://www.sciencedirect.com/science/article/pii/S0378434700006034},
author = {Laurie {Van Heukelem} and Crystal S Thomas},
keywords = {Carotenoids, Chlorophylls},
abstract = {We used chromatography modeling software to assist in HPLC method development, with the goal of enhancing separations through the exclusive use of gradient time and column temperature. We surveyed nine stationary phases for their utility in pigment purification and natural sample analysis. For purification, a complex algal matrix was separated on an efficient monomeric column, from which partially purified fractions were collected and purified on polymeric columns that exaggerated resolution between pigments of interest. Additionally, we feature an HPLC method that is simple, fast, demonstrates excellent transferability and is ideal for quantitative analysis of pigments in dilute natural water samples.}
}

@article{Fahy2005ALipids,
    title = {{A comprehensive classification system for lipids}},
    year = {2005},
    journal = {Journal of Lipid Research},
    author = {Fahy, Eoin and Subramaniam, Shankar and Brown, H. Alex and Glass, Christopher K. and Merrill, Alfred H. and Murphy, Robert C. and Raetz, Christian R.H. and Russell, David W. and Seyama, Yousuke and Shaw, Walter and Shimizu, Takao and Spener, Friedrich and Van Meer, Gerrit and VanNieuwenhze, Michael S. and White, Stephen H. and Witztum, Joseph L. and Dennis, Edward A.},
    number = {5},
    month = {5},
    pages = {839--861},
    volume = {46},
    publisher = {American Society for Biochemistry and Molecular Biology},
    url = {http://www.jlr.org/content/46/5/839.full http://www.jlr.org/content/46/5/839 http://www.jlr.org/content/46/5/839.abstract /content/46/5/839},
    doi = {10.1194/jlr.E400004-JLR200},
    issn = {00222275},
    pmid = {15722563},
    keywords = {Chemical representation, Fatty acyls, Glycerolipids, Glycerophospholipids, Informatics, Lipidomics, Nomenclature, Polyketides, Prenol lipids, Saccharolipids, Sphingolipids, Sterol lipids}
}

@article{JAHN1995157,
title = {Extraction of extracellular polymeric substances (EPS) from biofilms using a cation exchange resin},
journal = {Water Science and Technology},
volume = {32},
number = {8},
pages = {157-164},
year = {1995},
note = {Biofilm Structure, Growth and Dynamics},
issn = {0273-1223},
doi = {https://doi.org/10.1016/0273-1223(96)00020-0},
url = {https://www.sciencedirect.com/science/article/pii/0273122396000200},
author = {A. Jahn and P.H. Nielsen},
keywords = {Biofilm, cation exchange resin, cell lysis, extracellular polymeric substances, extraction},
abstract = {A cation exchange resin was used to extract water soluble EPS material from different sewer biofilms and a biofilm monoculture of P.putida. The extraction method seemed to be especially suitable for extracting humic substances and proteins from biofilms but also carbohydrate could be extracted, depending on the nature of the sample. Extraction conditions must be well defined because the extraction efficiency was dependent on stirring intensity, amount of cation exchange resin and extraction time. Up to 20\% of the total biofilm protein was found in the DOWEX-extractable fraction. No significant cell-lysis up to 2 hours of extraction time could be detected by acridine orange (AO) direct counts. The same was true when applying the extraction method to a batch culture of P.putida. However, the number of viable cells in the batch culture as recorded by plate count technique was reduced to 28\% after 30 mm of extraction. In conclusion, proteins and humic substances can be extracted from biofilm EPS in considerable amounts and should be investigated more in detail when analysing the properties of the EPS matrix of a biofilm.}
}

@article{hubas_nanoscale_2017,
	title = {A nanoscale study of carbon and nitrogen fluxes in mats of purple sulfur bacteria: {Implications} for carbon cycling at the surface of coastal sediments},
	volume = {8},
	issn = {1664302X},
	doi = {10.3389/fmicb.2017.01995},
	number = {OCT},
	journal = {Frontiers in Microbiology},
	author = {Hubas, C and Boeuf, D and Jesus, B and Thiney, N and Bozec, Y and Jeanthon, C},
	year = {2017},
	keywords = {Chromatiaceae, Compound specific isotope analysis (CSIA), Fatty acid methyl ester (FAME), Microbial mats, NanoSIMS, Stable isotope probing (SIP), Thiohalocapsa},
}

@article{bligh_rapid_1959,
	title = {A {Rapid} {Method} of {Total} {Lipid} {Extraction} and {Purification}},
	volume = {37},
	issn = {0576-5544},
	url = {https://cdnsciencepub.com/doi/10.1139/o59-099},
	doi = {10.1139/o59-099},
	number = {8},
	journal = {Canadian Journal of Biochemistry and Physiology},
	author = {Bligh, E G and Dyer, W J},
	year = {1959},
	pages = {911--917},
}

@article{gaubert-boussarie_untargeted_2020,
	title = {An untargeted metabolomic approach for microphytobenthic biofilms in intertidal mudflats},
	volume = {7},
	issn = {2296-7745},
	doi = {10.3389/FMARS.2020.00250},
	journal = {Frontiers in Marine Science},
	author = {Gaubert-Boussarie, Julie and Prado, Soizic and Hubas, Cédric},
	year = {2020},
	note = {Publisher: Frontiers},
	keywords = {Biofilms, Diatoms, Metabolomics, microphytobenthos, mudflats},
	pages = {250},
}

@article{decho_autoinducers_2009,
	title = {Autoinducers extracted from microbial mats reveal a surprising diversity of {N}-acylhomoserine lactones ({AHLs}) and abundance changes that may relate to diel {pH}.},
	volume = {11},
	issn = {1462-2920},
	url = {http://www.ncbi.nlm.nih.gov/pubmed/19196272},
	doi = {10.1111/j.1462-2920.2008.01780.x},
	number = {2},
	journal = {Environmental microbiology},
	author = {Decho, Alan W and Visscher, Pieter T and Ferry, John and Kawaguchi, Tomohiro and He, Lijian and Przekop, Kristen M and Norman, R Sean and Reid, R Pamela},
	year = {2009},
	pmid = {19196272},
	keywords = {Acyl-Butyrolactones, Acyl-Butyrolactones: classification, Acyl-Butyrolactones: isolation \{\&\} purification, Acyl-Butyrolactones: metabolism, Bacteria, Bacteria: chemistry, Bacteria: metabolism, Darkness, Geologic Sediments, Geologic Sediments: microbiology, Hydrogen-Ion Concentration, Light, Mass Spectrometry},
	pages = {409--420},
}

@incollection{cescutti_bacterial_2010,
	title = {Bacterial {Capsular} {Polysaccharides} and {Exopolysaccharides}},
	isbn = {978-0-12-374546-0},
	url = {https://doi.org/10.1016/B978-0-12-374546-0.00006-7},
	booktitle = {Microbial {Glycobiology}},
	publisher = {Elsevier Inc.},
	author = {Cescutti, Paola},
	month = jan,
	year = {2010},
	pages = {93--108},
}

@article{decho_bacterial_1990,
	title = {Bacterial exopolymer utilization by a harpacticoid copepod: a methodology and results},
	volume = {35},
	doi = {10.4319/lo.1990.35.5.1039},
	number = {5},
	journal = {Limnology and Oceanography},
	author = {Decho, A W and Moriarty, D J W},
	year = {1990},
	pages = {1039--1049},
}

@article{camilli_bacterial_2006,
	title = {Bacterial {Small}-{Molecule} {Signaling} {Pathways}},
	volume = {311},
	issn = {0036-8075},
	url = {http://www.ncbi.nlm.nih.gov/pubmed/16497924 http://www.pubmedcentral.nih.gov/articlerender.fcgi?artid=PMC2776824 http://www.sciencemag.org/cgi/doi/10.1126/science.1121357},
	doi = {10.1126/science.1121357},
	number = {5764},
	journal = {Science},
	author = {Camilli, A and Bassler, Bonnie L},
	year = {2006},
	pmid = {16497924},
	pages = {1113--1116},
}

@article{solano_biofilm_2014,
	title = {Biofilm dispersion and quorum sensing},
	volume = {18},
	issn = {13695274},
	url = {http://linkinghub.elsevier.com/retrieve/pii/S1369527414000290},
	doi = {10.1016/j.mib.2014.02.008},
	number = {1},
	journal = {Current Opinion in Microbiology},
	author = {Solano, Cristina and Echeverz, Maite and Lasa, Iñigo},
	year = {2014},
	pmid = {24657330},
	note = {ISBN: 1879-0364 (Electronic)\${\textbackslash}textbackslashbackslash\{{\textbackslash}textbackslash\$\}r1369-5274 (Linking)},
	pages = {96--104},
}

@article{bellinger_biofilm_2005,
	title = {Biofilm polymers: relationship between carbohydrate biopolymers from estuarine mudflats and unialgal cultures of benthic diatoms},
	volume = {38},
	issn = {0948-3055},
	url = {http://www.int-res.com/abstracts/ame/v38/n2/p169-180/},
	doi = {10.3354/ame038169},
	number = {2},
	journal = {Aquatic Microbial Ecology},
	author = {Bellinger, B J and Abdullahi, A S and Gretz, M R and Underwood, G J C},
	month = feb,
	year = {2005},
	note = {Publisher: Inter-Research},
	keywords = {Biofilms, Biopolymers, Diatoms, EPS, Fractionation, Microphytobenthos, Monosaccharide distribution, Uronic acids},
	pages = {169--180},
}

@article{costerton_biofilms_1994,
	title = {Biofilms, the {Customized} {Microniche}},
	volume = {176},
	doi = {10.1128/jb.176.8.2137-2142.1994},
	number = {8},
	journal = {Journal of Bacteriology},
	author = {Costerton, J W and Lewandowski, Z and De Beer, D and Caldwell, D and Korber, D and James, G},
	year = {1994},
	pages = {2137--2142},
}

@article{branda_biofilms_2005,
	title = {Biofilms: the matrix revisited.},
	volume = {13},
	issn = {0966-842X},
	url = {http://www.ncbi.nlm.nih.gov/pubmed/15639628},
	doi = {10.1016/j.tim.2004.11.006},
	number = {1},
	journal = {Trends in microbiology},
	author = {Branda, Steven S and Vik, Shild and Friedman, Lisa and Kolter, Roberto},
	month = jan,
	year = {2005},
	pmid = {15639628},
	keywords = {Bacterial, Bacterial: biosynthesis, Bacterial: physiology, Biofilms, Biofilms: growth \{\&\} development, Extracellular Matrix, Extracellular Matrix: chemistry, Extracellular Matrix: physiology, Fimbriae, Gene Expression Regulation, Polysaccharides},
	pages = {20--26},
}

@article{leinweber_capsules_2015,
	title = {Capsules of the diatom {Achnanthidium} minutissimum arise from fibrillar precursors and foster attachment of bacteria.},
	volume = {3},
	issn = {2167-8359},
	url = {http://www.pubmedcentral.nih.gov/articlerender.fcgi?artid=4380156%7B&%7Dtool=pmcentrez%7B&%7Drendertype=abstract},
	doi = {10.7717/peerj.858},
	journal = {PeerJ},
	author = {Leinweber, Katrin and Kroth, Peter G},
	month = jan,
	year = {2015},
	pmid = {25834772},
	pages = {e858},
}

@article{blair_carbon_1985,
	title = {Carbon isotopic fractionation in heterotrophic microbial metabolism.},
	volume = {50},
	issn = {00992240},
	url = {/pmc/articles/PMC291782/?report=abstract https://www.ncbi.nlm.nih.gov/pmc/articles/PMC291782/ http://www.pubmedcentral.nih.gov/articlerender.fcgi?artid=291782%7B&%7Dtool=pmcentrez%7B&%7Drendertype=abstract},
	doi = {10.1128/aem.50.4.996-1001.1985},
	number = {4},
	journal = {Applied and Environmental Microbiology},
	author = {Blair, N and Leu, A and Muñoz, E and Olsen, J and Kwong, E and Des Marais, D and Munoz, E and Olsen, J and Kwong, E and Des Marais, D},
	month = oct,
	year = {1985},
	pmid = {2867741},
	note = {Publisher: American Society for Microbiology (ASM)},
	keywords = {Acetates, Acetates: metabolism, Aspartic Acid, Aspartic Acid: metabolism, Carbon Isotopes, Citrates, Citrates: metabolism, Escherichia coli, Escherichia coli: metabolism, Glucose, Glucose: metabolism, Glutamates, Glutamates: metabolism, Glutamic Acid, Isotope Labeling, Isotope Labeling: methods, Kinetics},
	pages = {996--1001},
}

@article{summons_carbon_1994,
	title = {Carbon isotopic fractionation in lipids from methanotrophic bacteria: {Relevance} for interpretation of the geochemical record of biomarkers},
	volume = {58},
	issn = {00167037},
	doi = {10.1016/0016-7037(94)90119-8},
	number = {13},
	journal = {Geochimica et Cosmochimica Acta},
	author = {Summons, Roger E and Jahnke, Linda L and Roksandic, Zarko},
	month = jul,
	year = {1994},
	pmid = {11540111},
	note = {Publisher: Pergamon},
	pages = {2853--2863},
}

@article{vyssotski_chemical_2015,
	title = {Chemical synthesis and gas chromatographic behaviour of γ-stearidonic (18:4n-6) acid},
	volume = {92},
	issn = {0003021X},
	url = {https://link.springer.com/article/10.1007/s11746-014-2588-x},
	doi = {10.1007/s11746-014-2588-x},
	number = {3},
	journal = {JAOCS, Journal of the American Oil Chemists' Society},
	author = {Vyssotski, Mikhail and Lagutin, Kirill and MacKenzie, Andrew and Itabashi, Yutaka},
	month = mar,
	year = {2015},
	note = {Publisher: Springer Verlag},
	keywords = {GC, GC-MS, NMR, Octadecapentaenoic, Octadecatetraenoic, Stearidonic, Δ3 PUFA},
	pages = {383--391},
}

@article{vallet_chemically-mediated_2018,
	title = {Chemically-{Mediated} {Interactions} {Between} {Macroalgae}, {Their} {Fungal} {Endophytes}, and {Protistan} {Pathogens}},
	volume = {9},
	issn = {1664-302X},
	doi = {10.3389/fmicb.2018.03161},
	journal = {Frontiers in Microbiology},
	author = {Vallet, Marine and Strittmatter, Martina and Murúa, Pedro and Lacoste, Sandrine and Dupont, Joëlle and Hubas, Cedric and Genta-Jouve, Gregory and Gachon, Claire M M and Kim, Gwang Hoon and Prado, Soizic},
	month = dec,
	year = {2018},
	note = {Publisher: Frontiers Media SA},
}

@article{dubois_colorimetric_1956,
	title = {Colorimetric {Method} for {Determination} of {Sugars} and {Related} {Substances}},
	volume = {28},
	doi = {10.1021/ac60111a017},
	number = {3},
	journal = {Analytical Chemistry},
	author = {Dubois, M and Gilles, K A and Hamilton, J K and Rebers, P A and Smith, F},
	year = {1956},
	pages = {350--356},
}

@article{dijkman_composition_2010,
	title = {Composition and heterogeneity of the microbial community in a coastal microbial mat as revealed by the analysis of pigments and phospholipid-derived fatty acids},
	volume = {63},
	issn = {13851101},
	url = {http://linkinghub.elsevier.com/retrieve/pii/S1385110109000987},
	doi = {10.1016/j.seares.2009.10.002},
	number = {1},
	journal = {Journal of Sea Research},
	author = {Dijkman, Nicole a. and Boschker, Henricus T S and Stal, Lucas J and Kromkamp, Jacco C},
	month = jan,
	year = {2010},
	note = {Publisher: Elsevier B.V.},
	pages = {62--70},
}

@article{oakes_composition_2010,
	title = {Composition, production, and loss of carbohydrates in subtropical shallow subtidal sandy sediments: {Rapid} processing and long-term retention revealed by {13C}-labeling},
	volume = {55},
	issn = {1939-5590},
	url = {http://dx.doi.org/10.4319/lo.2010.55.5.2126 http://doi.wiley.com/10.4319/lo.2010.55.5.2126},
	doi = {10.4319/lo.2010.55.5.2126},
	number = {5},
	journal = {Limnology and Oceanography},
	author = {Oakes, Joanne M and Eyre, Bradley D and Middelburg, Jack J and Boschker, Henricus T S},
	month = sep,
	year = {2010},
	note = {Publisher: John Wiley \& Sons, Ltd},
	pages = {2126--2138},
}

@inproceedings{tolhurst_diatom_2003,
	title = {Diatom migration and sediment armouring - {An} example from the {Tagus} {Estuary}, {Portugal}},
	volume = {503},
	url = {www.sediments.co.uk},
	doi = {10.1023/B:HYDR.0000008474.33782.8d},
	booktitle = {Hydrobiologia},
	publisher = {Springer},
	author = {Tolhurst, T J and Jesus, B and Brotas, V and Paterson, D M},
	month = aug,
	year = {2003},
	note = {ISSN: 00188158
Issue: 1},
	keywords = {Chlorophyll a, Diatom migration, Microphytobenthos, Sediment stability},
	pages = {183--193},
}

@article{haynes_diatom-derived_2007,
	title = {Diatom-derived carbohydrates as factors affecting bacterial community composition in estuarine sediments},
	volume = {73},
	issn = {0099-2240},
	url = {http://www.ncbi.nlm.nih.gov/pubmed/17675437 http://www.pubmedcentral.nih.gov/articlerender.fcgi?artid=PMC2075028 http://aem.asm.org/cgi/doi/10.1128/AEM.00551-07},
	doi = {10.1128/AEM.00551-07},
	number = {19},
	journal = {Applied and Environmental Microbiology},
	author = {Haynes, K and Hofmann, T A and Smith, C J and Ball, A S and Underwood, G J C and Osborn, A M},
	month = oct,
	year = {2007},
	pmid = {17675437},
	pages = {6112--6124},
}

@article{bohorquez_different_2017,
	title = {Different types of diatom-derived extracellular polymeric substances drive changes in heterotrophic bacterial communities from intertidal sediments},
	volume = {8},
	issn = {1664302X},
	url = {/pmc/articles/PMC5326797/?report=abstract https://www.ncbi.nlm.nih.gov/pmc/articles/PMC5326797/},
	doi = {10.3389/fmicb.2017.00245},
	number = {FEB},
	journal = {Frontiers in Microbiology},
	author = {Bohórquez, Julio and McGenity, Terry J and Papaspyrou, Sokratis and García-Robledo, Emilio and Corzo, Alfonso and Underwood, Graham J C},
	month = feb,
	year = {2017},
	note = {Publisher: Frontiers Research Foundation},
	keywords = {Degradation rate, EPS, G-model, Intertidal sediment, Microbial community, Microphytobenthos, Nutrient regeneration, Pyrosequencing},
	pages = {245},
}

@article{hanlon_dynamics_2006,
	title = {Dynamics of extracellular polymeric substance ({EPS}) production and loss in an estuarine, diatom-dominated, microalgal biofilm over a tidal emersio-immersion period},
	volume = {51},
	issn = {00243590},
	url = {http://doi.wiley.com/10.4319/lo.2006.51.1.0079},
	doi = {10.4319/lo.2006.51.1.0079},
	number = {1},
	journal = {Limnology and Oceanography},
	author = {Hanlon, A R M and Bellinger, B and Haynes, K and Xiao, G and Hofmann, T A and Gretz, M R and Ball, A S and Osborn, A M and Underwood, G J C},
	month = jan,
	year = {2006},
	pages = {79--93},
}

@article{cartaxana_effects_2008,
	title = {Effects of two motility inhibitors on the photosynthetic activity of the diatoms {Cylindrotheca} closterium and {Pleurosigma} angulatum},
	volume = {23},
	issn = {21598347},
	url = {https://www.tandfonline.com/doi/abs/10.1080/0269249X.2008.9705737},
	doi = {10.1080/0269249X.2008.9705737},
	number = {1},
	journal = {Diatom Research},
	author = {Cartaxana, Paulo and Brotas, Vanda and Serôdio, João},
	year = {2008},
	note = {Publisher: Taylor \& Francis Group},
	pages = {65--74},
}

@article{findlay_efficacy_1989,
	title = {Efficacy of phospholipid analysis in determining microbial biomass in sediments},
	volume = {55},
	issn = {00992240},
	url = {/pmc/articles/PMC203186/?report=abstract https://www.ncbi.nlm.nih.gov/pmc/articles/PMC203186/},
	doi = {10.1128/aem.55.11.2888-2893.1989},
	number = {11},
	journal = {Applied and Environmental Microbiology},
	author = {Findlay, R H and King, G M and Watling, L},
	year = {1989},
	pmid = {16348051},
	note = {Publisher: American Society for Microbiology (ASM)},
	pages = {2888--2893},
}

@article{dunstan_essential_1994,
	title = {Essential {Polyunsaturated} {Fatty} {Acids} from 14 {Species} of {Diatoms} ({Bacillariophycaee})},
	volume = {35},
	doi = {10.1016/S0031-9422(00)90525-9},
	number = {1},
	journal = {Phytochemistry},
	author = {Dunstan, Graeme A and Volkman, John K and Barrett, Stephanie M and Leroi, Jeannie-marie and Jeffrey, S W},
	year = {1994},
	keywords = {biotechnology, diatoms, index-bacillariophycae, polyunsaturated fatty acids},
	pages = {155--161},
}

@article{yi_exploring_2017,
	title = {Exploring valuable lipids in diatoms},
	volume = {4},
	url = {https://www.frontiersin.org/articles/10.3389/fmars.2017.00017/full},
	doi = {10.3389/fmars.2017.00017},
	number = {17},
	author = {Yi, Zhiqian and Xu, Maonian and Di, Xiaxia and Brynjolfsson, Sigurdur and Fu, Weiqi},
	month = jan,
	year = {2017},
	note = {ISSN: 22967745
Issue: JAN
Pages: 17
Publication Title: Frontiers in Marine Science
Volume: 4
DOI:},
	keywords = {Diatoms, Fatty acids, Lipid production, Oxylipin, Value-added products},
}

@article{veuger_fate_2012,
	title = {Fate of microbial nitrogen, carbon, hydrolysable amino acids, monosaccharides, and fatty acids in sediment},
	volume = {83},
	issn = {00167037},
	doi = {10.1016/j.gca.2011.12.016},
	journal = {Geochimica et Cosmochimica Acta},
	author = {Veuger, Bart and van Oevelen, Dick and Middelburg, Jack J},
	month = apr,
	year = {2012},
	note = {Publisher: Pergamon},
	pages = {217--233},
}

@article{jonasdottir_fatty_2019,
	title = {Fatty acid profiles and production in marine phytoplankton},
	volume = {17},
	doi = {10.3390/md17030151},
	number = {3},
	author = {Jónasdóttir, Sigrún Huld},
	month = mar,
	year = {2019},
	doi = {10.3390/md17030151},
	pmid = {30836652},
	note = {ISSN: 16603397
Issue: 3
Pages: 151
Publication Title: Marine Drugs
Volume: 17},
	keywords = {DHA, Environmental effects, EPA, Fatty acid synthesis, Fatty acids, Marine phytoplankton, PUFA synthesis, SDA},
	pages = {151},
}

@article{de_carvalho_fatty_2014,
	title = {Fatty acids as a tool to understand microbial diversity and their role in food webs of mediterranean temporary ponds},
	volume = {19},
	issn = {14203049},
	url = {/pmc/articles/PMC6271346/?report=abstract https://www.ncbi.nlm.nih.gov/pmc/articles/PMC6271346/},
	doi = {10.3390/molecules19055570},
	number = {5},
	journal = {Molecules},
	author = {De Carvalho, Carla C C R C R and Caramujo, Maria José},
	year = {2014},
	pmid = {24786844},
	note = {ISBN: 1420-3049
Publisher: Molecular Diversity Preservation International},
	keywords = {Autotrophic biomass, Bacteria, Bacterial identification, Heterotrophic biomass, PLFA, Trophic level},
	pages = {5570--5598},
}

@article{sahu_fatty_2013,
	title = {Fatty acids as biomarkers of microalgae},
	volume = {89},
	issn = {00319422},
	doi = {10.1016/j.phytochem.2013.02.001},
	journal = {Phytochemistry},
	author = {Sahu, Abhishek and Pancha, Imran and Jain, Deepti and Paliwal, Chetan and Ghosh, Tonmoy and Patidar, Shailesh and Bhattacharya, Sourish and Mishra, Sandhya},
	month = may,
	year = {2013},
	pmid = {23453131},
	note = {Publisher: Pergamon},
	keywords = {Algorithmic hierarchical clustering, Biomarker, Chemotaxonomy, Fatty acids, Microalgae, Nutrition, Principal component analysis},
	pages = {53--58},
}

@article{kharlamenko_fatty_1995,
	title = {Fatty acids as markers of food sources in a shallow-water hydrothermal ecosystem ({Kraternaya} {Bight}, {Yankich} {Island}, {Kurile} {Islands})},
	volume = {120},
	doi = {10.3354/meps120231},
	journal = {Marine Ecology-Progress Series},
	author = {Kharlamenko, V I and Zhukova, N V and Khotimchenko, S V and Svetashev, V I and Kamenev, G M},
	year = {1995},
	keywords = {acid, biomarkers, food web - fatty, v e n t},
	pages = {231--241},
}

@article{meziane_fatty_2000,
	title = {Fatty acids as tracers of organic matter in the sediment and food web of a mangrove/intertidal flat ecosystem, {Okinawa}, {Japan}},
	volume = {200},
	doi = {10.3354/meps200049},
	journal = {Marine Ecology Progress Series},
	author = {Meziane, T and Tsuchiya, M},
	year = {2000},
	pages = {49--57},
}

@article{perry_fatty_1979,
	title = {Fatty acids of bacterial origin in contemporary marine sediments},
	volume = {43},
	issn = {00167037},
	doi = {10.1016/0016-7037(79)90020-6},
	number = {11},
	journal = {Geochimica et Cosmochimica Acta},
	author = {Perry, G J and Volkman, J K and Johns, R B and Bavor, H J},
	month = nov,
	year = {1979},
	note = {Publisher: Pergamon},
	pages = {1715--1725},
}

@article{le_bris_hyperspectral_2016,
	title = {Hyperspectral remote sensing of wild oyster reefs},
	volume = {172},
	issn = {0272-7714},
	doi = {10.1016/J.ECSS.2016.01.039},
	journal = {Estuarine, Coastal and Shelf Science},
	author = {Le Bris, Anthony and Rosa, Philippe and Lerouxel, Astrid and Cognie, Bruno and Gernez, Pierre and Launeau, Patrick and Robin, Marc and Barillé, Laurent},
	month = apr,
	year = {2016},
	note = {Publisher: Academic Press},
	keywords = {Benthos, Crassostrea gigas, Hyperspectral, Oyster reefs, Remote sensing, Tidal flats},
	pages = {1--12},
}

@article{kaneda_iso-_1991,
	title = {Iso- and {Anteiso}-{Fatty} {Acids} in {Bacteria} : {Biosynthesis} , {Function} , and {Taxonomic} {Significance}},
	volume = {55},
	doi = {10.1128/mr.55.2.288-302.1991},
	number = {2},
	journal = {Microbiological Reviews},
	author = {Kaneda, Toshi},
	year = {1991},
	pages = {288--302},
}

@article{pegu_isolation_2020,
	title = {Isolation and characterization of a-l-rhamnosidase producing bacterium, {Agrococcus} sp. bkd37, from a warehouse soil and partial optimization of its culture conditions},
	volume = {22},
	issn = {0972-0979},
	doi = {10.5958/0974-4517.2020.00028.2},
	number = {3},
	journal = {Applied Biological Research},
	author = {Pegu, Bhaba Kumar and Kardong, Devid and Chetia, Pankaj and Chutia, Jitu and Gogoi, Dip Kumar},
	year = {2020},
	note = {Publisher: Diva Enterprises Private Limited},
	pages = {203--214},
}

@article{vallet_laminariales_2020,
	title = {Laminariales host does impact lipid temperature trajectories of the fungal endophyte paradendryphiella salina ({Sutherland}.)},
	volume = {18},
	issn = {16603397},
	url = {www.mdpi.com/journal/marinedrugs},
	doi = {10.3390/MD18080379},
	number = {8},
	journal = {Marine Drugs},
	author = {Vallet, Marine and Meziane, Tarik and Thiney, Najet and Prado, Soizic and Hubas, Cedric},
	month = jul,
	year = {2020},
	pmid = {32708010},
	note = {Publisher: MDPI AG},
	keywords = {Fatty acids, Fungal endophytes, Laminariales, Paradendryphiella salina},
	pages = {379},
}

@article{combe_mapping_2005,
	title = {Mapping microphytobenthos biomass by non-linear inversion of visible-infrared hyperspectral images},
	volume = {98},
	issn = {0034-4257},
	doi = {10.1016/J.RSE.2005.07.010},
	number = {4},
	journal = {Remote Sensing of Environment},
	author = {Combe, Jean Philippe and Launeau, Patrick and Carrère, Véronique and Despan, Daniela and Méléder, Vona and Barillé, Laurent and Sotin, Christophe},
	month = oct,
	year = {2005},
	note = {Publisher: Elsevier},
	keywords = {Biomass, Continuum, Imaging spectroscopy, Mapping, Microphytobenthos, Modified Gaussian Model, Reflectance spectroscopy, Spectral library},
	pages = {371--387},
}

@article{paul_methane-oxidizing_2017,
	title = {Methane-oxidizing bacteria shunt carbon to microbial mats at a marine hydrocarbon seep},
	volume = {8},
	issn = {1664302X},
	doi = {10.3389/FMICB.2017.00186/BIBTEX},
	number = {FEB},
	journal = {Frontiers in Microbiology},
	author = {Paul, Blair G and Ding, Haibing and Bagby, Sarah C and Kellermann, Matthias Y and Redmond, Molly C and Andersen, Gary L and Valentine, David L},
	month = feb,
	year = {2017},
	note = {Publisher: Frontiers Research Foundation},
	keywords = {16S rRNA gene, Intact polar lipids (IPL), Methanotrophs, Microbial mats, Stable isotope probing, Sulfide-oxidizing bacteria},
	pages = {186},
}

@article{uhlig_methane-oxidizing_2018,
	title = {Methane-oxidizing seawater microbial communities from an {Arctic} shelf},
	volume = {15},
	issn = {17264189},
	doi = {10.5194/BG-15-3311-2018},
	number = {11},
	journal = {Biogeosciences},
	author = {Uhlig, Christiane and Kirkpatrick, John B and D'Hondt, Steven and Loose, Brice},
	month = jun,
	year = {2018},
	note = {Publisher: Copernicus GmbH},
	pages = {3311--3329},
}

@article{decho_microbial_2000,
	title = {Microbial biofilms in intertidal systems : an overview},
	volume = {20},
	doi = {10.1016/S0278-4343(00)00022-4},
	number = {10-11},
	journal = {Continental Shelf Research},
	author = {Decho, A W},
	year = {2000},
	pages = {1257--1273},
}

@article{guerrero_microbial_2002,
	title = {Microbial mats and the search for minimal ecosystems.},
	volume = {5},
	issn = {1139-6709},
	url = {http://www.ncbi.nlm.nih.gov/pubmed/12497183},
	doi = {10.1007/s10123-002-0094-8},
	number = {4},
	journal = {International microbiology : the official journal of the Spanish Society for Microbiology},
	author = {Guerrero, R and Piqueras, M and Berlanga, M},
	year = {2002},
	pmid = {12497183},
	keywords = {Biofilms, Biofilms: growth \{\&\} development, Biological, Biomass, Carbon, Carbon: metabolism, Cyanobacteria, Cyanobacteria: growth \{\&\} development, Cyanobacteria: ultrastructure, Ecosystem, Environmental Microbiology, Models, Sulfides, Sulfides: metabolism},
	pages = {177--188},
}

@article{taylor_microphytobenthic_2013,
	title = {Microphytobenthic extracellular polymeric substances ( {EPS} ) in intertidal sediments fuel both generalist and specialist {EPS}-degrading bacteria},
	volume = {58},
	doi = {10.4319/lo.2013.58.4.1463},
	number = {4},
	journal = {Limnology and Oceanography},
	author = {Taylor, Joe D and Mckew, Boyd A and Kuhl, Alison and Mcgenity, Terry J and Underwood, Graham J C},
	year = {2013},
	pages = {1463--1480},
}

@article{kazemipour_microphytobenthos_2012,
	title = {Microphytobenthos biomass mapping using the optical model of diatom biofilms: {Application} to hyperspectral images of {Bourgneuf} {Bay}},
	volume = {127},
	issn = {0034-4257},
	doi = {10.1016/J.RSE.2012.08.016},
	journal = {Remote Sensing of Environment},
	author = {Kazemipour, Farzaneh and Launeau, Patrick and Méléder, Vona},
	month = dec,
	year = {2012},
	note = {Publisher: Elsevier},
	keywords = {Biofilm, Biomass, Bourgneuf Bay, Chl a, Diatom, Euglenid, Hyperspectral remote sensing, Microphytobenthos},
	pages = {1--13},
}

@article{takahashi_optimising_2009,
	title = {Optimising extraction of extracellular polymeric substances ({EPS}) from benthic diatoms: comparison of the efficiency of six {EPS} extraction methods},
	volume = {60},
	issn = {1323-1650},
	url = {http://www.publish.csiro.au/?paper=MF08258},
	doi = {10.1071/MF08258},
	number = {12},
	journal = {Marine and Freshwater Research},
	author = {Takahashi, Eri and Ledauphin, Jérôme and Goux, Didier and Orvain, Francis},
	year = {2009},
	note = {Publisher: CSIRO publishing},
	keywords = {Biofilm, Cell lysis, Dowex, GCMS analysis, Immuno staining, Immunostaining, Intertidal sediments, Navicula jeffreyi, Protein, Transmission electron microscopy},
	pages = {1\{{\textbackslash}textbackslashtextendash\}10},
}

@article{passarelli_organisms_2014,
	title = {Organisms as cooperative ecosystem engineers in intertidal flats},
	volume = {92},
	issn = {13851101},
	doi = {10.1016/j.seares.2013.07.010},
	journal = {Journal of Sea Research},
	author = {Passarelli, C and Olivier, F and Paterson, D M M and Meziane, T and Hubas, C},
	year = {2014},
	keywords = {[Biogenic structure, Biogenic structure, Cooperative ecosystem enginee, Cooperative ecosystem engineers, Habitat cascade, Sediment stability, Tidal flats},
}

@article{decho_overview_2010,
	title = {Overview of biopolymer-induced mineralization: {What} goes on in biofilms?},
	volume = {36},
	issn = {09258574},
	url = {http://linkinghub.elsevier.com/retrieve/pii/S0925857409001128},
	doi = {10.1016/j.ecoleng.2009.01.003},
	number = {2},
	journal = {Ecological Engineering},
	author = {Decho, Alan W},
	year = {2010},
	note = {Publisher: Elsevier B.V.},
	pages = {137--144},
}

@article{cartaxana_physiological_2011,
	title = {Physiological versus behavioral photoprotection in intertidal epipelic and epipsammic benthic diatom communities},
	volume = {405},
	issn = {00220981},
	url = {http://linkinghub.elsevier.com/retrieve/pii/S0022098111002425},
	doi = {10.1016/j.jembe.2011.05.027},
	number = {1-2},
	journal = {Journal of Experimental Marine Biology and Ecology},
	author = {Cartaxana, P and Ruivo, M and Hubas, C and Davidson, I and Serôdio, J and Jesus, B},
	year = {2011},
	keywords = {Diatom, Microphytobenthos, Migration, Photoprotection, Xanthophyll cycle},
	pages = {120--127},
}

@article{hubas_proliferation_2013,
	title = {Proliferation of purple sulphur bacteria at the sediment surface affects intertidal mat diversity and functionality},
	volume = {8},
	issn = {19326203},
	doi = {10.1371/journal.pone.0082329},
	number = {12},
	journal = {PLoS ONE},
	author = {Hubas, C and Jesus, B and Ruivo, M and Meziane, T and Thiney, N and Davoult, D and Spilmont, N and Paterson, D M M and Jeanthon, C},
	year = {2013},
}

@article{lowry_protein_1951,
	title = {Protein measurement with the folin phenol reagent},
	volume = {193},
	issn = {00219258},
	url = {https://linkinghub.elsevier.com/retrieve/pii/S0021925819524516},
	doi = {10.1016/S0021-9258(19)52451-6},
	number = {1},
	journal = {Journal of Biological Chemistry},
	author = {Lowry, OliverH. and Rosebrough, NiraJ. and Farr, A Lewis and Randall, RoseJ.},
	month = nov,
	year = {1951},
	pages = {265--275},
}

@article{passarelli_seasonal_2015,
	title = {Seasonal variations of the composition of microbial biofilms in sandy tidal flats: {Focus} of fatty acids, pigments and exopolymers},
	volume = {153},
	issn = {02727714},
	url = {http://www.sciencedirect.com/science/article/pii/S0272771414003412},
	doi = {10.1016/j.ecss.2014.11.013},
	number = {0},
	journal = {Estuarine, Coastal and Shelf Science},
	author = {Passarelli, C and Meziane, T and Thiney, N and Boeuf, D and Jesus, B and Ruivo, M and Jeanthon, C and Hubas, C},
	year = {2015},
	keywords = {[Biofilms, Biofilms, Biogenic structures, EPS, Fatty acids, Monosaccharide composition, Pigments},
	pages = {29--37},
}

@article{boon_significance_1977,
	title = {Significance and taxonomic value of iso and anteiso monoenoic fatty acids and branded beta-hydroxy acids in {Desulfovibrio} desulfuricans},
	volume = {129},
	url = {https://journals.asm.org/doi/10.1128/jb.129.3.1183-1191.1977},
	doi = {10.1128/jb.129.3.1183-1191.1977},
	number = {3},
	journal = {Journal of Bacteriology},
	author = {Boon, Jaap J and Leeuw, J W De and v.D. Hoek, G J and Vosjan, H J and de Leeuw, J W and v. d. Hoek, G J and Vosjan, J H},
	year = {1977},
	pages = {1183--1191},
}

@article{bellinger_significance_2009,
	title = {Significance of diatom-derived polymers in carbon flow dynamics within estuarine biofilms determined through isotopic enrichment},
	volume = {55},
	url = {http://www.int-res.com/abstracts/ame/v55/n2/p169-187/},
	doi = {10.3354/ame01287},
	number = {2},
	journal = {Aquatic Microbial Ecology},
	author = {Bellinger, B J and Underwood, Graham J C and Ziegler, S E and Gretz, Michael R},
	year = {2009},
	pages = {169--187},
}

@article{meleder_spatio-temporal_2005,
	title = {Spatio-temporal changes in microphytobenthos structure analysed by pigment composition in a macrotidal flat ({Bourgneuf} {Bay}, {France})},
	volume = {297},
	issn = {0171-8630},
	url = {http://www.int-res.com/abstracts/meps/v297/p83-99/},
	doi = {10.3354/meps297083},
	journal = {Marine Ecology Progress Series},
	author = {Méléder, V and Barillé, L and Rincé, Y and Morançais, M and Rosa, P and Gaudin, P},
	year = {2005},
	keywords = {diatoms, high-pressure liquid chromatography, microphytobenthos, pigments},
	pages = {83--99},
}

@article{meleder_spatiotemporal_2007,
	title = {Spatiotemporal changes in microphytobenthos assemblages in a macrotidal flat ({Bourgneuf} {Bay}, {France})},
	volume = {43},
	issn = {00223646},
	doi = {10.1111/J.1529-8817.2007.00423.X},
	number = {6},
	journal = {Journal of Phycology},
	author = {Méléder, Vona and Rincé, Yves and Barillé, Laurent and Gaudin, Pierre and Rosa, Philippe},
	month = dec,
	year = {2007},
	keywords = {Assemblage, Biomass, Life-form, macrotidal flat, Microphytobenthos},
	pages = {1177--1190},
}

@article{gladyshev_stable_2012,
	title = {Stable isotope composition of fatty acids in organisms of different trophic levels in the {Yenisei} {River}.},
	volume = {7},
	issn = {1932-6203},
	url = {http://www.pubmedcentral.nih.gov/articlerender.fcgi?artid=3314690%7B&%7Dtool=pmcentrez%7B&%7Drendertype=abstract},
	doi = {10.1371/journal.pone.0034059},
	number = {3},
	journal = {PloS one},
	author = {Gladyshev, Michail I and Sushchik, Nadezhda N and Kalachova, Galina S and Makhutova, Olesia N},
	month = jan,
	year = {2012},
	pmid = {22470513},
	keywords = {Animals, Carbon Isotopes, Carbon Isotopes: chemistry, Fatty Acids, Fatty Acids: chemistry, Food Chain, Larva, Larva: metabolism, Microalgae, Microalgae: metabolism, Rivers},
	pages = {e34059},
}

@article{remize_study_2020,
	title = {Study of synthesis pathways of the essential polyunsaturated fatty acid 20:5n-3 in the diatom chaetoceros muelleri using13 c-isotope labeling},
	volume = {10},
	issn = {2218273X},
	url = {www.mdpi.com/journal/biomolecules},
	doi = {10.3390/biom10050797},
	number = {5},
	journal = {Biomolecules},
	author = {Remize, Marine and Planchon, Frédéric and Loh, Ai Ning and Le Grand, Fabienne and Bideau, Antoine and Le Goic, Nelly and Fleury, Elodie and Miner, Philippe and Corvaisier, Rudolph and Volety, Aswani and Soudant, Philippe},
	month = may,
	year = {2020},
	pmid = {32455747},
	note = {Publisher: MDPI AG},
	keywords = {20:5n-3 (EPA), Acyl-editing mechanism, Chaetoceros muelleri, Compound-specific isotope analysis, Diatom, Synthesis pathway},
	pages = {797},
}

@article{wotton_essential_2004,
	title = {The essential role of exopolymers ({EPS}) in aquatic systems},
	volume = {42},
	url = {https://doi.org/10.1201/9780203507810},
	doi = {10.1201/9780203507810},
	journal = {Oceanography and Marine Biology: an Annual Review},
	author = {Wotton, R S},
	year = {2004},
	pages = {57--94},
}

@article{middelburg_fate_2000,
	title = {The fate of intertidal microphytobenthos carbon: {An} in situ {13C}-labeling study},
	volume = {45},
	issn = {00243590},
	doi = {10.4319/lo.2000.45.6.1224},
	number = {6},
	journal = {Limnology and Oceanography},
	author = {Middelburg, Jack J and Barranguet, Christiane and Boschker, Henricus T S and Herman, Peter M J and Moens, Tom and Heip, Carlo H R},
	year = {2000},
	note = {ISBN: 0024-3590},
	pages = {1224--1234},
}

@article{underwood_importance_2003,
	series = {Advances in {Botanical} {Research}},
	title = {The importance of extracellular carbohydrate productionby marine epipelic diatoms},
	volume = {40},
	url = {https://doi.org/10.1016/S0065-2296(05)40005-1},
	doi = {10.1016/S0065-2296(05)40005-1},
	author = {Underwood, Graham J C and Paterson, David M},
	year = {2003},
	doi = {10.1016/s0065-2296(05)40005-1},
	note = {ISSN: 0065-2296},
	pages = {183--240},
}

@article{lubarsky_stabilisation_2010,
	title = {The {Stabilisation} {Potential} of {Individual} and {Mixed} {Assemblages} of {Natural} {Bacteria} and {Microalgae}},
	volume = {5},
	issn = {19326203},
	url = {http://www.pubmedcentral.nih.gov/articlerender.fcgi?artid=2970557%7B&%7Dtool=pmcentrez%7B&%7Drendertype=abstract},
	doi = {10.1371/journal.pone.0013794},
	number = {11},
	journal = {PLoS ONE},
	author = {Lubarsky, H V and Hubas, C and Chocholek, M and Larson, F and Manz, W and Paterson, D M and Gerbersdorf, S U},
	month = jan,
	year = {2010},
	pmid = {21072186},
	keywords = {Bacteria, Bacteria: genetics, Bacteria: growth \{\&\} development, Bacteria: metabolism, Biomass, Biopolymers, Biopolymers: metabolism, Chlorophyll, Chlorophyll: metabolism, Diatoms, Diatoms: growth \{\&\} development, Diatoms: metabolism, Diatoms: ultrastructure, Ecosystem, Electron, Flow Cytometry, Fluorescence, Geologic Sediments, Geologic Sediments: microbiology, In Situ Hybridization, Microalgae, Microalgae: growth \{\&\} development, Microalgae: metabolism, Microalgae: ultrastructure, Microscopy, Pheophytins, Pheophytins: metabolism, Scanning},
	pages = {e13794},
}

@article{leu_ultraviolet_2007,
	title = {Ultraviolet radiation negatively affects growth but not food quality of arctic diatoms},
	volume = {52},
	issn = {00243590},
	url = {http://doi.wiley.com/10.4319/lo.2007.52.2.0787},
	doi = {10.4319/lo.2007.52.2.0787},
	number = {2},
	journal = {Limnology and Oceanography},
	author = {Leu, Eva and Falk-Petersen, Stig and Hessen, Dag O},
	month = mar,
	year = {2007},
	note = {Publisher: American Society of Limnology and Oceanography Inc.},
	pages = {787--797},
}
