## Supplementary figures and images for "Identification of microbial exopolymer producers in sandy and muddy intertidal sediments by compound-specific isotope analysis"

### Fig1.pdf

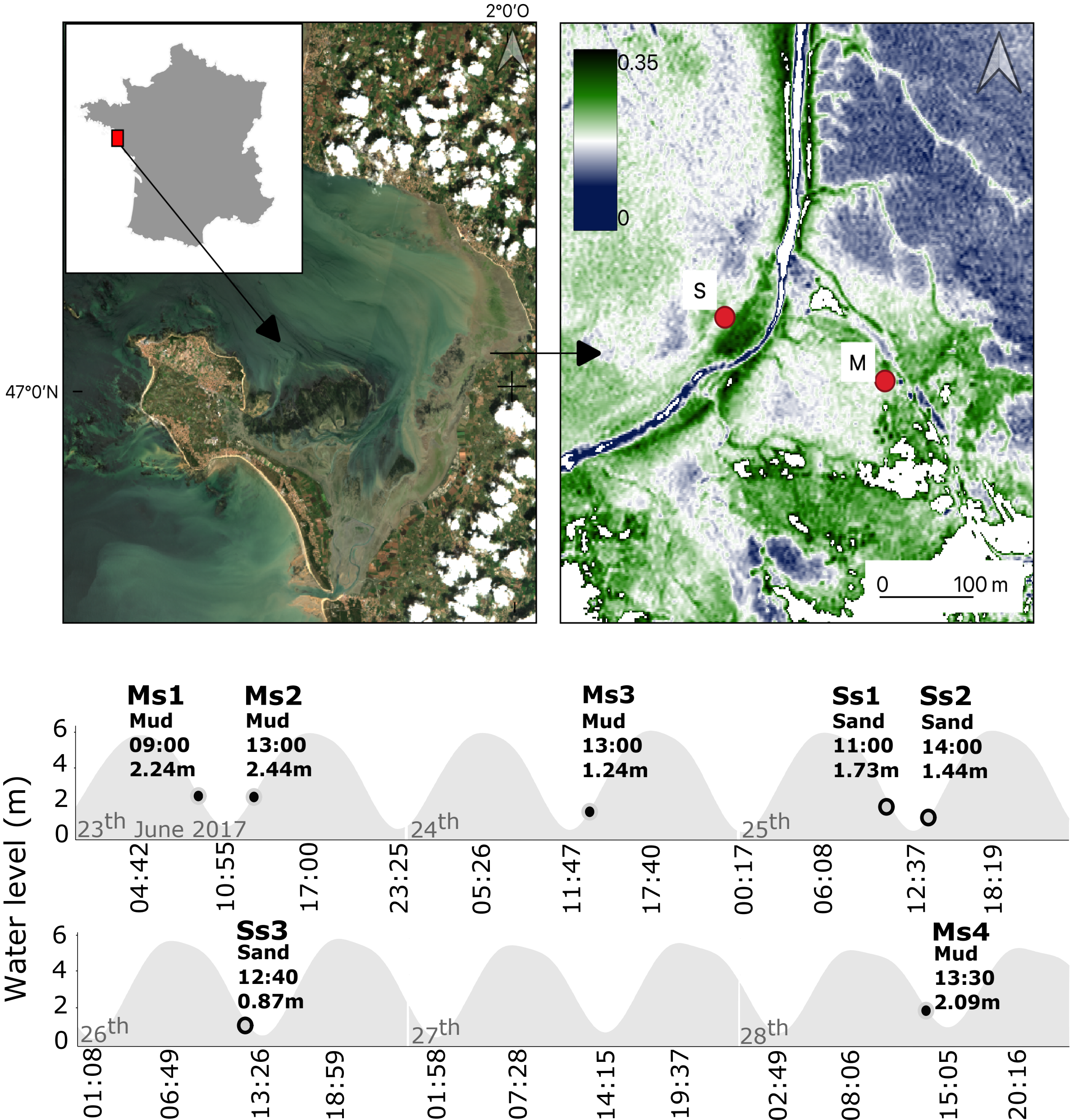

### Fig2.pdf

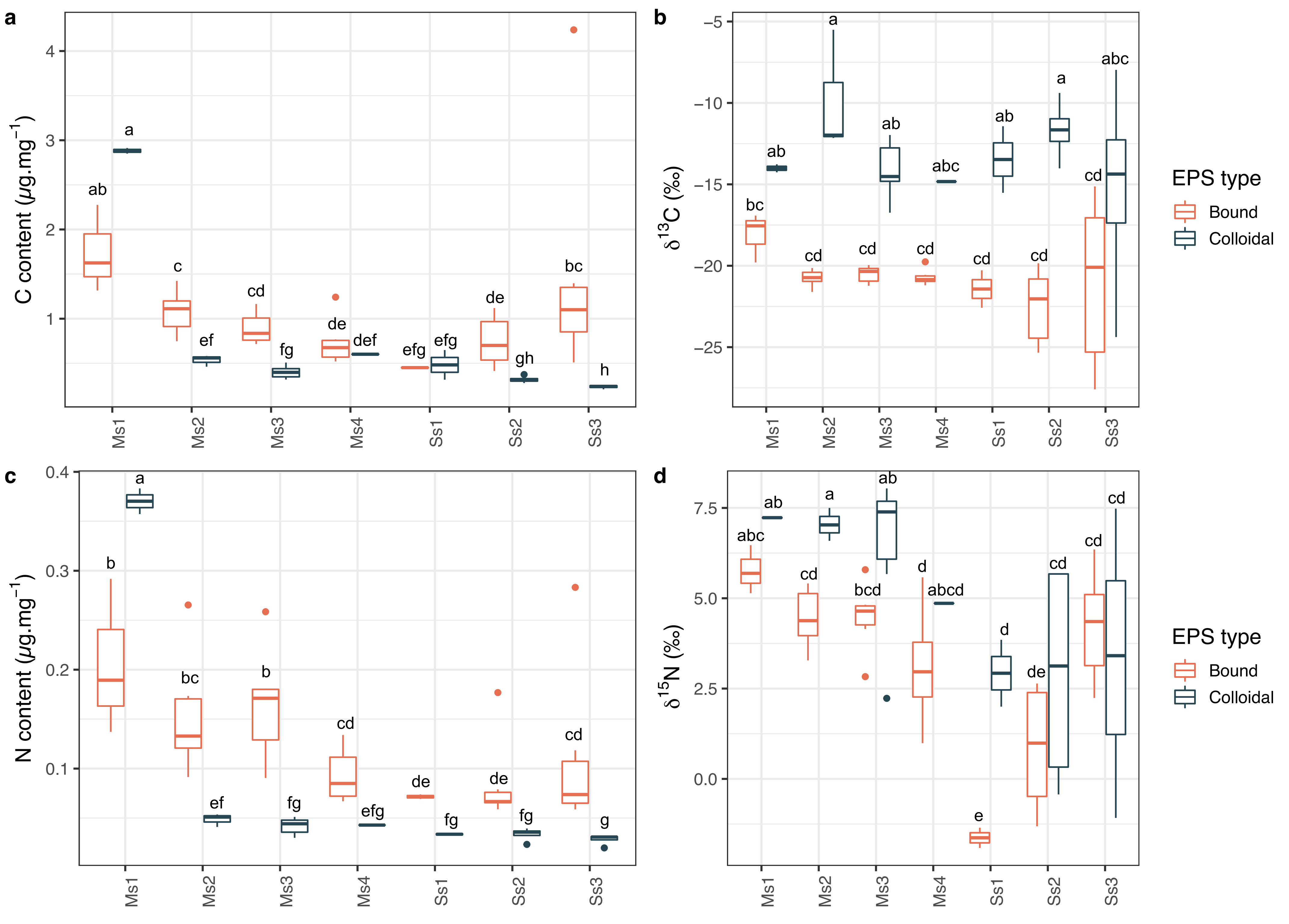

### Fig3.pdf

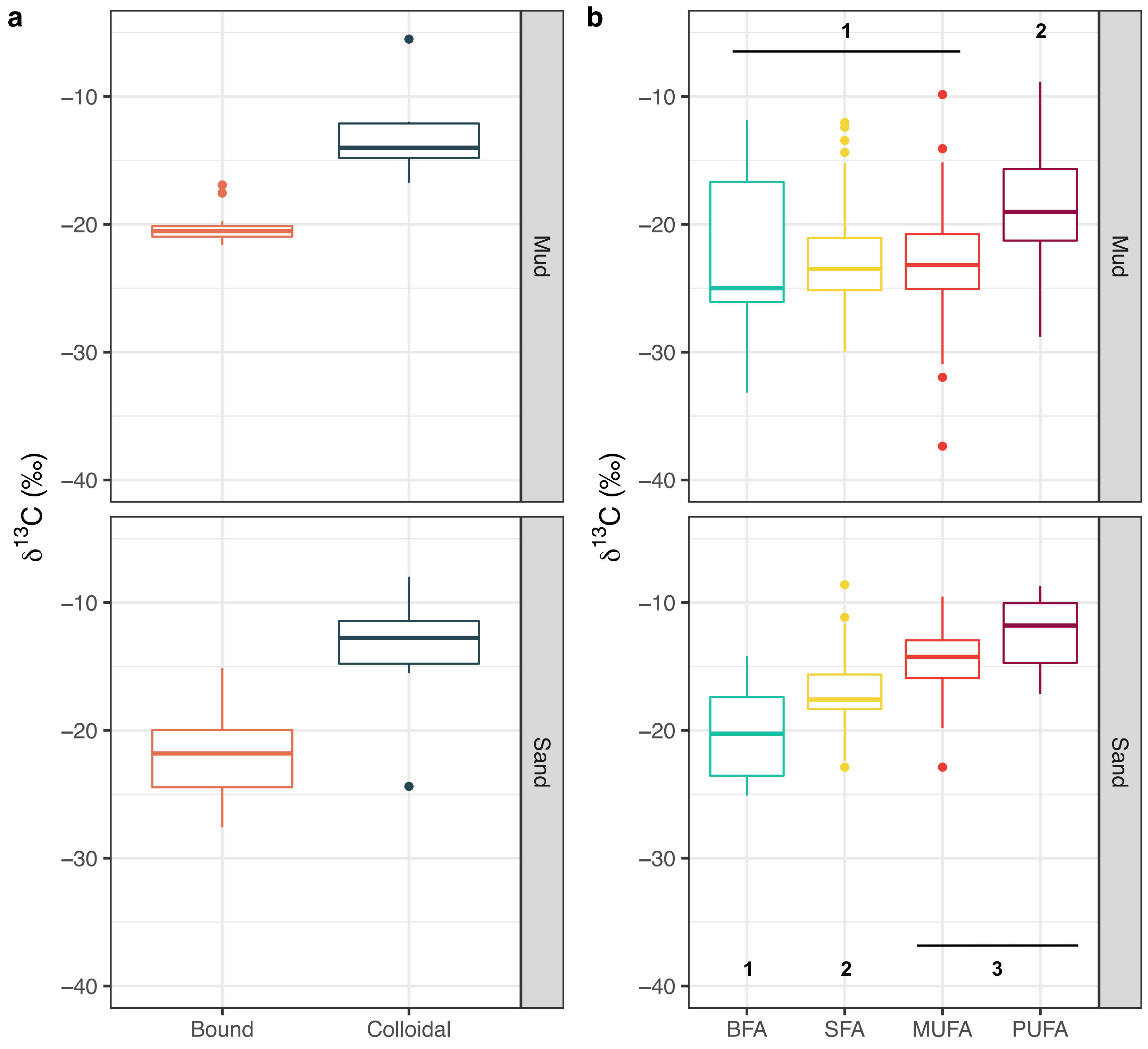

### Fig4.pdf

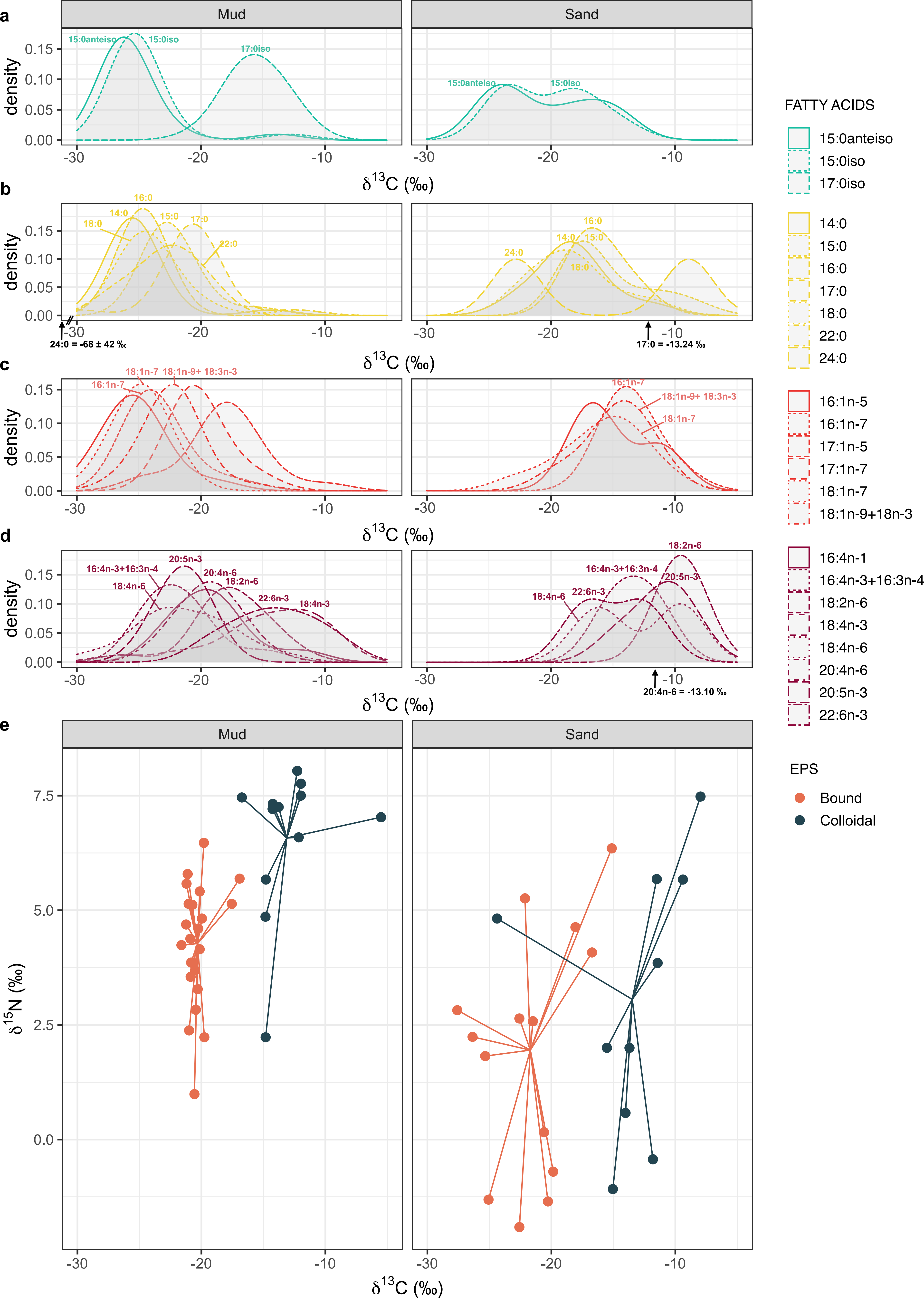

### Fig5.pdf

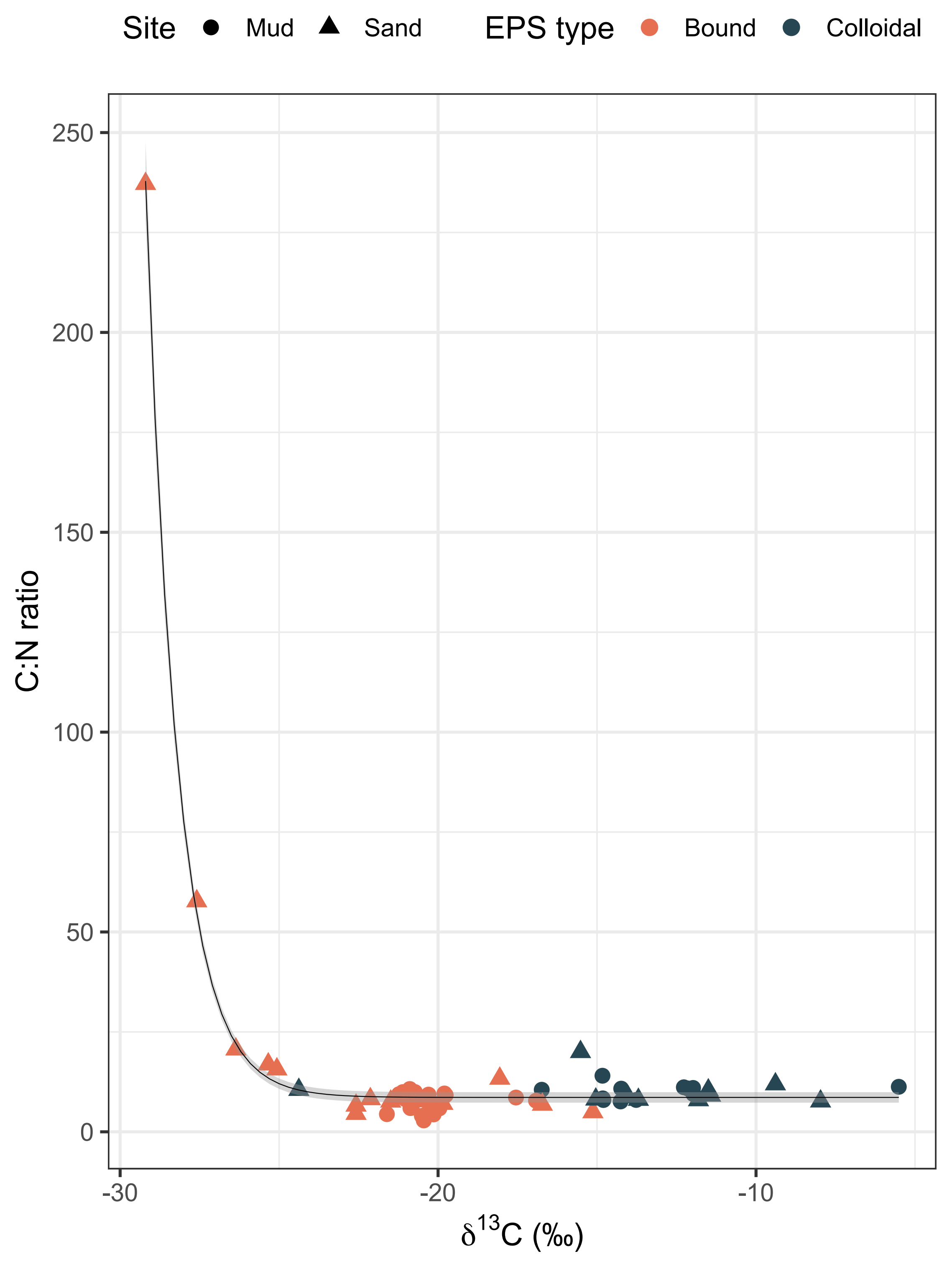

### FigSF1.pdf

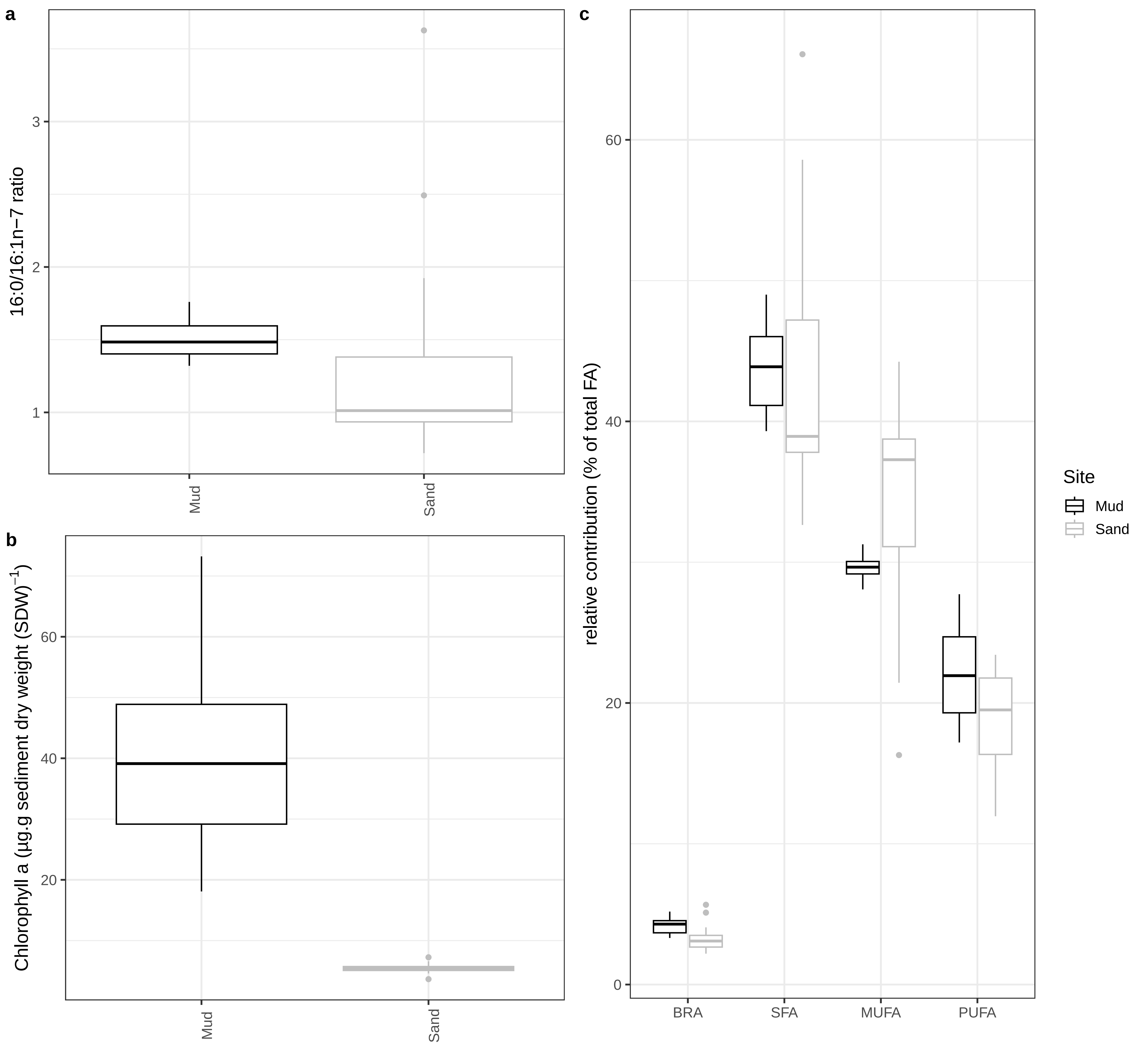

### FigSF2.pdf

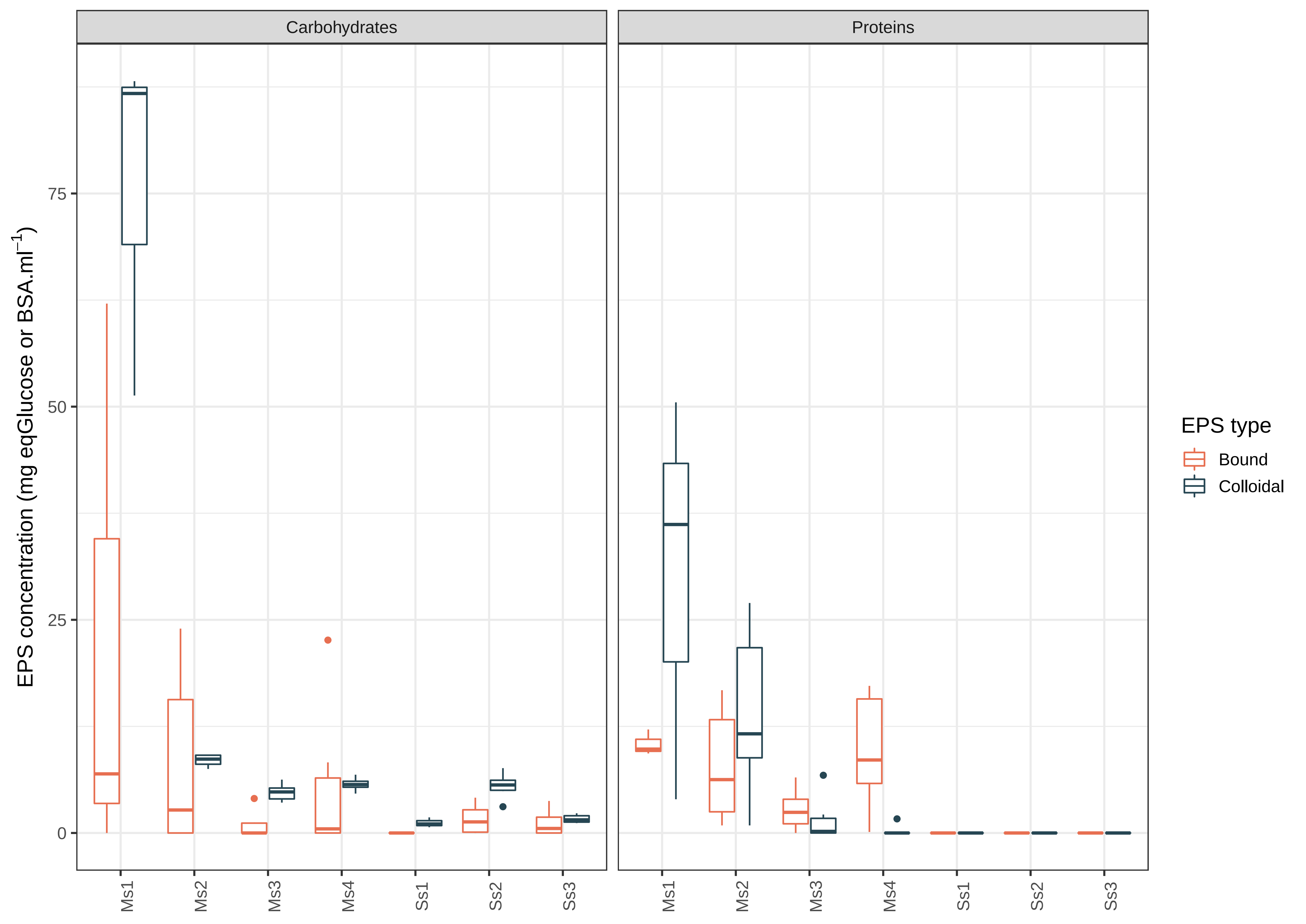
