## Supplementary material for "Identification of microbial exopolymer producers in sandy and muddy intertidal sediments by compound-specific isotope analysis": Revised_article_with_changes

**Extracellular polymeric substances (EPS) refer to a wide variety of high molecular weight molecules secreted outside the cell membrane by biofilm microorganisms. In the present study, EPS from marine microphytobenthic biofilms were extracted and their isotope ratios were analysed. A comparison of these ratios with the carbon isotope ratios of fatty acid biomarkers allowed the identification of the main EPS producers of two contrasting types of intertidal marine sediments. Our study reveals that ~~EPS production and degradation are supported by very different communities in muddy and sandy sediments and that~~ EPS sources are more diverse in sandy sediments than in muddy sediments. ~~In mud, bound EPS are mainly derived from diatoms, while colloidal EPS are the result of degradation of bound exopolymers by certain specialised bacteria. In sand, bound EPS are rather of bacterial or cyanobacterial origin and diatoms contribute mainly to colloidal EPS. These differences are thought to be related to differences in the functioning of the epipelic and epipsammic communities and in particular to the use of EPS either for motility or for cell attachment purposes.~~ We also found distinct patterns in the production and breakdown of EPS in sandy and muddy environments. The main difference observed was in how epipelic and epipsammic diatoms affected the chemistry of EPS, which had significant implications for the growth of bacteria specialized in utilizing EPS. These differences were likely linked to variations in the functioning of epipelic and epipsammic communities, specifically in how EPS was used either for motility or for cell attachment.**

In this study, we extracted colloidal and bound EPS from intertidal biofilms and analysed the natural stable isotope ratios (SIR) of carbon ($\delta^{13}$C) and nitrogen ($\delta^{15}$N). ~~Isotope ratios of EPS were compared to those of fatty acid biomarkers to determine which microorganisms were primarily responsible for the production of EPS in muddy and sandy sediments.~~ In order to identify the main contributors of EPS in muddy and sandy sediments, the SIR of EPS were compared to those of fatty acid biomarkers. These fatty acids are specific indicators of certain microorganisms, as their relative proportions vary distinctly across different organisms. For example, the major fatty acid in diatoms is 20:5n-3 (24–26). By examining the isotope ratios of EPS alongside these fatty acid biomarkers, the study aimed to determine the primary microorganisms responsible for EPS production.

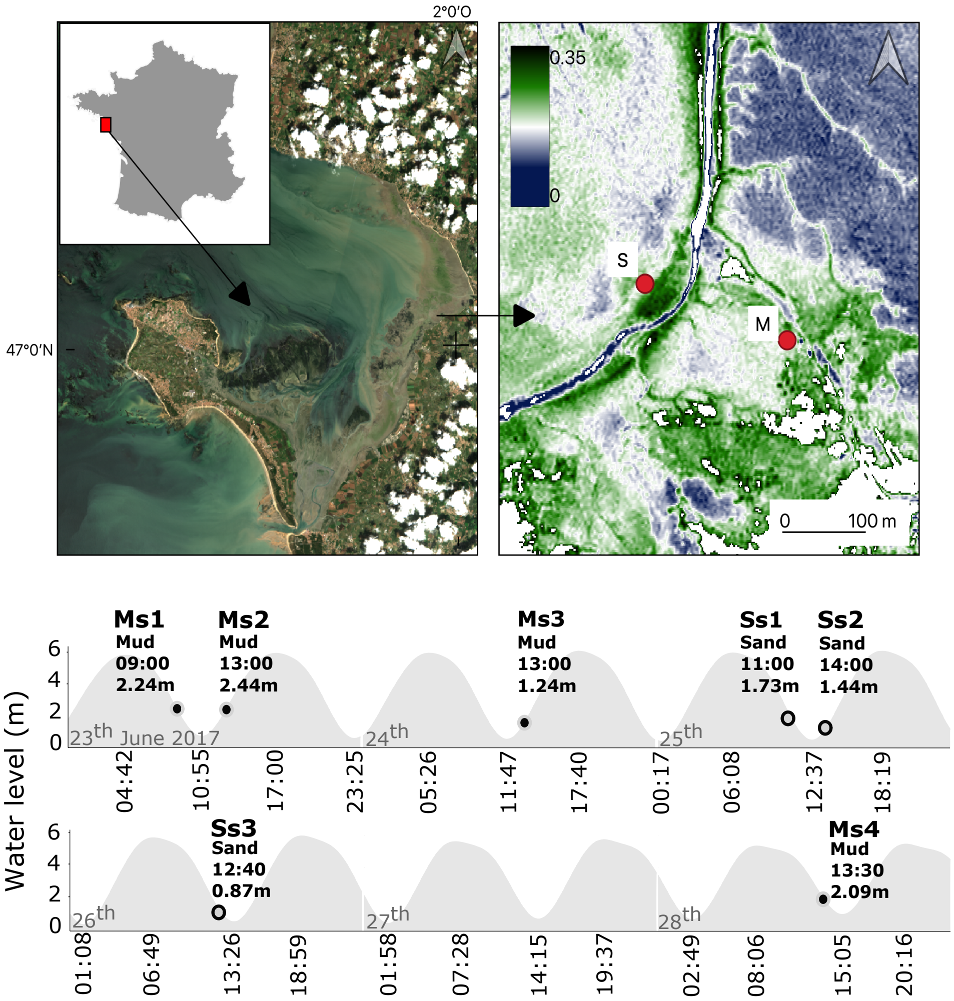


**Fig. 1.** Studied area. Top panel: Map of Bourgneuf Bay (France) and location of the sampling sites (Pleiades image acquired on the 2017/06/24 at 11:15 - UTC). left = true colors, right = Normalized Difference Vegetation Index (NDVI); Bottom panel: sampling occasions according to the tidal level at the study sites (data provided by the Naval Hydrographic and Oceanographic Service (SHOM) for coordinates: 047°06'00.0"N, 002°07' 00.0"W (Pornic). The first capital letter indicates the type of sediment (M=mud, S=sand), the other letters (s1 to s4) indicate the sampling point.

$$\delta^{13}\mathrm{C}_{FA} = \frac{(\delta^{13}\mathrm{C}_{FAME} - (1-f)\delta^{13}\mathrm{C}_{CH_3OH})}{f} \quad (1)$$

where $\delta^{13}\mathrm{C}_{FA}$ and $\delta^{13}\mathrm{C}_{FAME}$ (in ‰) are the isotopic composition of the free FA, and the FA methyl ester respectively, $f$ is the fractional carbon contribution of the free FA to the ester and $\delta^{13}\mathrm{C}_{CH_3OH}$ is the isotopic composition of the methanol derivatization reagent (–39.1 ‰).

~~**Fatty acid identification.** Identification of the samples was performed using a gas chromatograph coupled to mass spectrometer (GC–MS, Varian 450GC with Varian 220-MS). Compounds annotation was performed by comparing mass spectra with NIST 2017 library. Corresponding fatty acids are designated as X:Yn-Z, where X is the number of carbons, Y the number of double bonds and Z the position of the ultimate double bond from the terminal methyl (see (43) for additional information about naming convention).~~

**Table 1.** Comparison of Carbon and Nitrogen contents and isotopic ratios of colloidal and bound EPS at all sampling occasions using the van der Waerden (Normal Scores) non parametric test. df = degree of freedom. Results of the post-hoc test using the criterium Fisher's least significant difference (LSD) are shown in Fig. 2

| Variable | $\chi^2$ | df | p-value |
|---|---|---|---|
| Carbon content | 46.83024 | 13 | 1.03209e-05 |
| Nitrogen content | 27.364 | 14 | 6.74533e-06 |
| $\delta^{13}$C | 36.31005 | 13 | 0.00053 |
| $\delta^{15}$N | 35.02141 | 13 | 0.00084 |

We also examined the crossed impact of EPS and sediment type (i.e. BoundMud, BoundSand, ColloidalMud, and ColloidalSand). Our findings revealed significant differences between these levels in terms of C and N content and stable isotope ratios. For detailed information on these differences, please refer to table 2.

**EPS isotopic compositions.** At the muddy site, bound EPS were always $^{13}$C or $^{15}$N depleted in comparison to colloidal EPS at this site (Fig. 2b,d). At the sandy site, the same pattern is observed but both nitrogen and carbon stable isotope ratio showed a higher variability.

All sampling dates together, $\delta^{13}$C (Fig. 3a, top panel) and $\delta^{15}$N values were significantly different between bound and colloidal EPS at the muddy site (Permutation two Sample t-tests, $\delta^{13}$C: t = -10.678, p-value = 0.002; $\delta^{15}$N: t = -4.4325, p-value = 0.002).

At the sandy site, $\delta^{13}$C values were also significantly different (Fig. 3a, bottom panel) between bound and colloidal EPS (two Sample Student's t-test, t = -4.9474, df = 22, p-value = 5.984e-05) but $\delta^{15}$N was not significantly different (two Sample Student's t-test, t = -0.97547, df = 22, p-value = 0.3399).

All sampling dates grouped together $\delta^{13}$C between bound and colloidal EPS were thus always significantly different at both sites (Fig. 3a), indicating that these two fractions were from different EPS producers. Comparison with the literature is difficult as it is the first time that C and N natural stable isotopes ratio are reported on intertidal bound and colloidal EPS. The values reported in the literature for the main monosaccharides constituting the extracellular sugars are however in agreement with our results (i.e. a natural $\delta^{13}$C of -15 to -18 ‰) (23)

**Carbon isotope ratio of fatty acid classes.** In sandy sediment $\delta^{13}$C were significantly different between fatty acids classes ($F = 23.128, df1 = 3, df2 = 109, p = 1.16 \times 10^{-11}$) and showed a gradual $^{13}$C enrichment (Fig. 3b) from branched fatty acids (BFA) to mono- (MUFA) and polyunsaturated (PUFA) fatty acids. Such differences were not observed in the muddy site. In the mud, $\delta^{13}$C of BFA, saturated (SFA) and MUFA were not significantly different. Only PUFA showed a slightly higher mean $\delta^{13}$C (Permutation one-way Welch Anova followed by Tukey HSD posthoc test, $F = 33.588, p < 2.2 \times 10^{-16}$).

In comparison with similar ecosystems (i.e. intertidal muddy sediments), the isotope ratios of the main fatty acids are quite

**Table 2.** NEW TABLE:Comparison of Carbon and Nitrogen contents and isotopic ratios of colloidal and bound EPS in mud and sand. The sampling occasions for mud (Ms) and sand (Ms) were grouped together. The van der Waerden (Normal Scores) non-parametric test was used, with consideration of degrees of freedom (df). The post-hoc test results using Fisher's least significant difference (LSD) as the criterion are shown in the table

| | | Carbon content | Nitrogen content | $\delta^{13}$C | $\delta^{15}$N |
|---|---|---|---|---|---|
| Van der Waerden test statistics | $\chi^2$, df, p-value | 26.7, 3, <0.001 | 31.1, 3, <0.001 | 32.1, 3, <0.001 | 23.7, 3, <0.001 |
| Posthoc tests | Bound EPS - Mud | a | a | a | a |
| | Bound EPS - Sand | ab | b | a | b |
| | Colloidal EPS - Mud | b | b | b | bc |
| | Colloidal EPS - Sand | c | c | c | c |

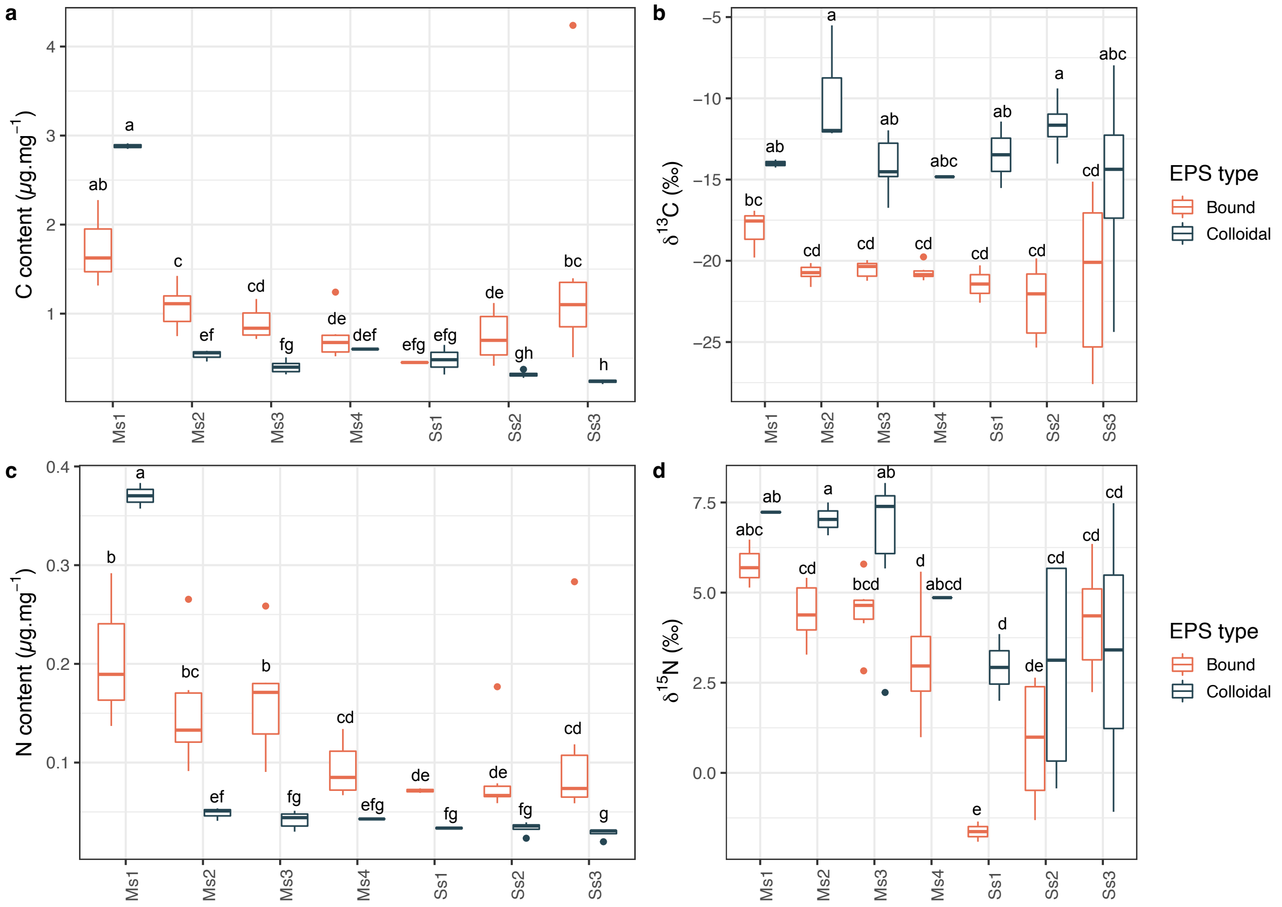


**Fig. 2.** Chemical composition of the EPS. **a**,**c**: Carbon (C) and Nitrogen (N) contents in $\mu$g per mg of freeze dried EPS. **b**,**d**: Carbon and Nitrogen stable isotope ratio ($\delta$ notation against atmospheric $N_2$ and Vienna PDB respectively) of the EPS. Colloidal EPS corresponded to loose, water-extractable exopolymers whereas bound EPS correspond to ion exchange resin-extractable exopolymers. Letters within the graph represent results of Fisher's least significant difference (LSD) post-hoc test. For the corresponding van der Waerden test, please see Table 1

consistent. Previous studies recorded $\delta^{13}$C ranging from -16 to -21‰ for branched, -14 to -26‰ for saturated, -13 to -22‰ for monounsaturated and -15 to -22‰ for polyunsaturated fatty acids (3, 22, 48). Taylor et al. (48) also showed that natural carbon isotope ratios were highly variable even over relatively short periods (i.e. 30h). These changes indicate that subtle modifications in the metabolic processes of carbon assimilation as well as interactions between microorganisms can take place over very short periods and could explain the variability of our $\delta^{13}$C values.

The tetracosanoic acid (SFA, 24:0) was excluded from the above mentioned analyses as it increased dramatically the variability because of extreme and unusually negative $\delta^{13}$C values indicative of a specific metabolism. The mean $\delta^{13}$C of 24:0 was $-66.89 \pm 35.84$‰ and $-59.24 \pm 71.82$‰ in the mud and sand respectively. It also sometimes showed a plurimodal distribution (as shown by density plots figure 4b) which indicate that 24:0 had likely varied microbial origins. This particular fatty acid was the only one to show extremely low $\delta^{13}$C values in line with the isotopic ratios generally found in methane-rich ecosystems for which direct links could be established between $\delta^{13}$C values and the presence of methane-oxidizers in bacterial communities (49, 50). It is indeed possible that the 24:0 originated from anaerobic bacteria related to the oxidation of methane or the sulphur cycle. The most negative $\delta^{13}$C values were recorded in highly re-

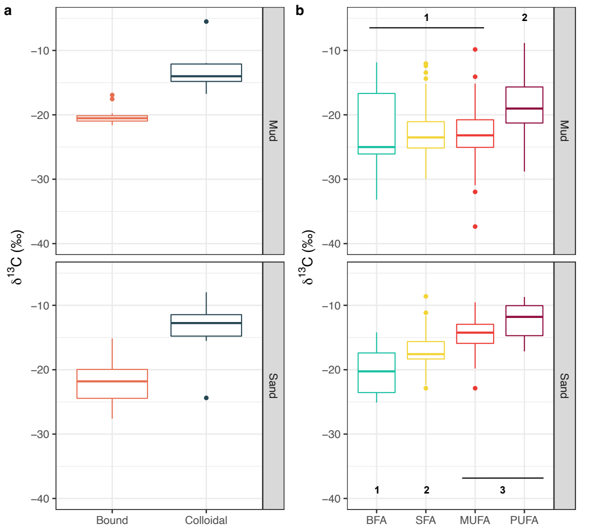


**Fig. 3.** Comparison of $\delta^{13}$C (corrected according to equation 1) between fatty acid classes and EPS fractions. **a**: bound and colloidal EPS were significantly different (t-tests, $p < 0.001$) at both sites. **b**: numbers indicate significantly different groups as evidenced by post-hoc tests. BFA=branched, SFA=saturated, MUFA=monounsaturated and PUFA=polyunsaturated fatty acids

duced muddy sediments.

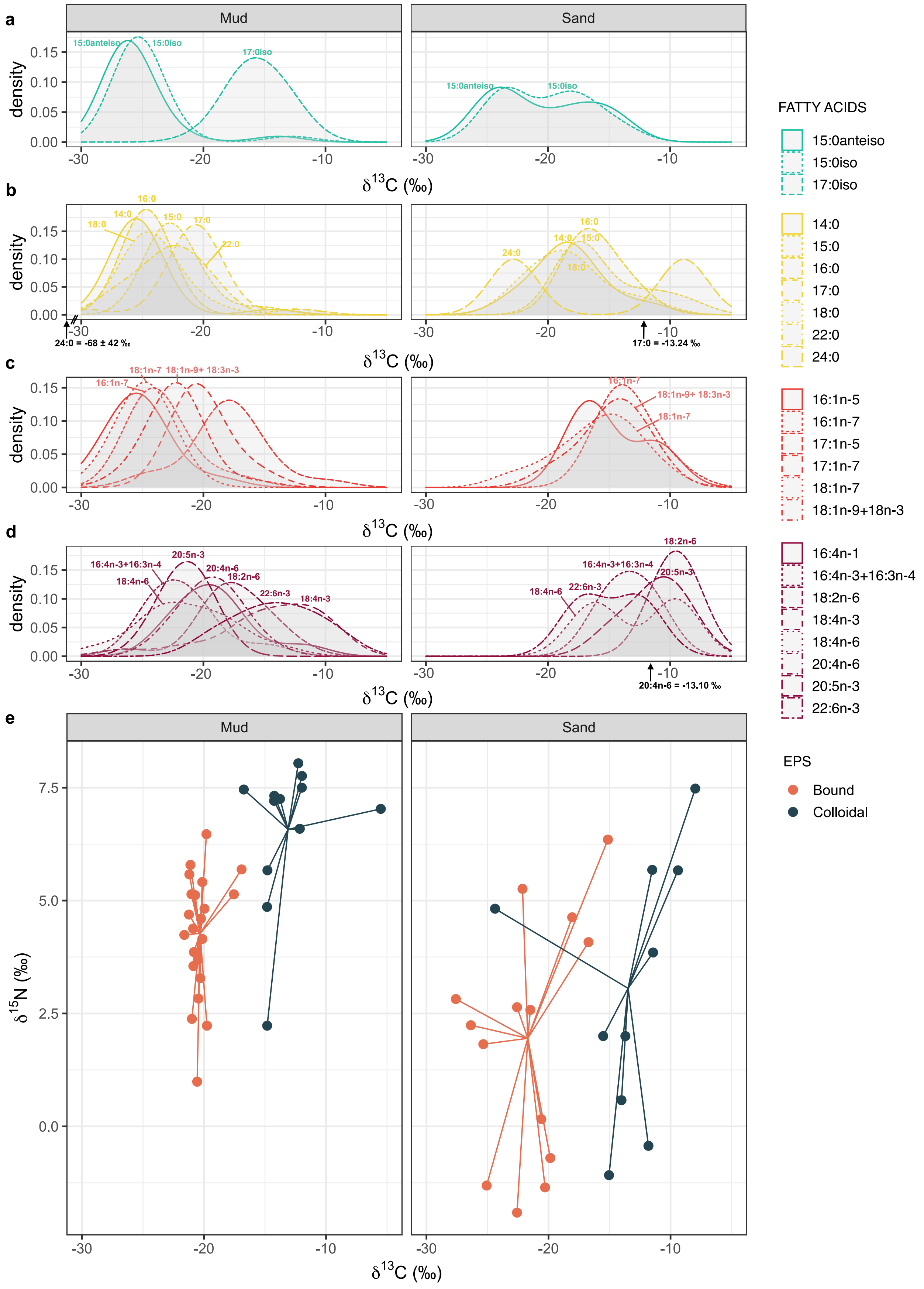


**Fig. 4.** C and N isotopic ratio of fatty acids and EPS fractions. **a-d**: Kernel density estimates of $\delta^{13}$C of fatty acid biomarkers (corrected according to equation 1). **e**: $\delta^{13}$C and $\delta^{15}$N biplots of bound and colloidal EPS. All sampling points were grouped together. In panels **a-d**, fatty acids are grouped by classes they belong to according to fig.3

only occurred in the bound fraction.

### CONFLICT OF INTEREST DISCLOSURE

The authors declare that they have no financial conflicts of interest in relation to the content of the article.

### AUTHOR CONTRIBUTIONS

Conceptualization: KS, BJ, CH ; Data Curation: CH ; Formal analysis: CH ; Funding acquisition: KS, BJ, CH ; Investigation: all authors ; Methodology: all authors ; Project administration: KS, BJ, CH ; Resources: all authors ; Software: CH ; Supervision: KS ; Validation: all authors ; Visualization: CH ; Writing – original draft: CH ; Writing – review & editing: CH, BJ, VM, KS.

### ACKNOWLEDGEMENTS

Some data were acquired on research platforms, which we would like to thank. CSIA data were acquired at Stable Isotope Platform of the European Institute for Marine Studies (IUEM, Brest, France). EA-IRMS data were acquired at the UC Davis stable isotope facility. The Pleiades image used in this study was ordered as part of the ISIS program of the French National Centre for Space Studies (CNES) and provided by DataTerra via the DINAMIS platform. The image is available on the Airbus Geostore catalogue (https://www.intelligence-airbusds.com/en/4871-geostore-ordering) with the following identifier: DS_PHR1B_201706241115350_FR1_PX_W003N47_1201_01276

### FUNDING

This study was supported by the BIO-Tide project (The role of microbial biodiversity in the functioning of marine tidal flat sediments), funded through the 2015–2016 BiodivERsA COFUND call for research proposals, with the national funders BelSPO (BRAIN-be contract BR/175/A1/BIO-Tide-BE), FWO (3G0H6816), ANR (ANR-16-EBI3-0008), and SNSF. We additionally acknowledge funding from FWO project G003820N. The post-doctoral grant of J. G-B was supported by the Regional Council of Brittany, SAD program and META-Tide projects.

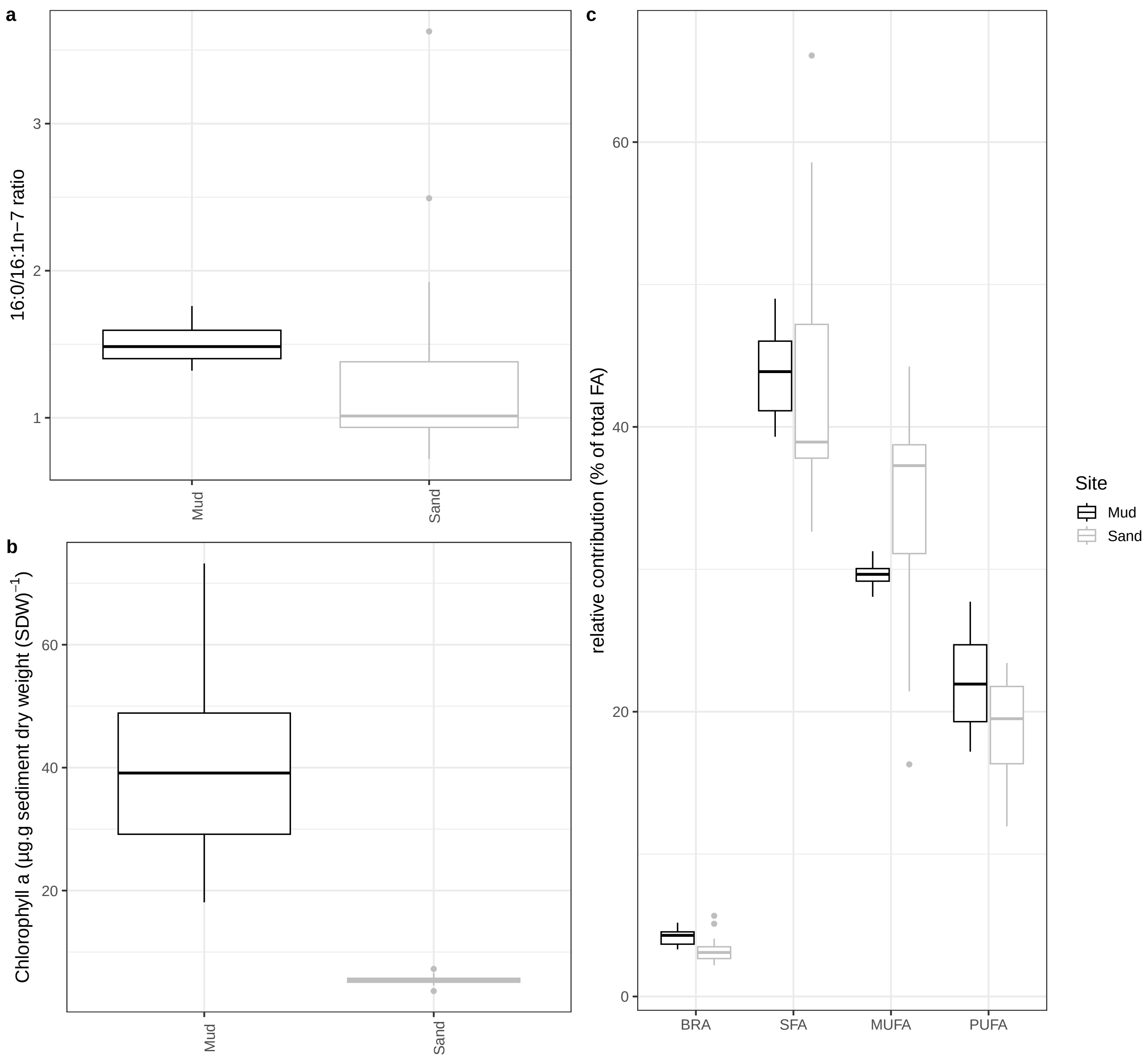


**Fig. SF1.** NEW FIGURE Comparison of biomass indicators and general composition of the microphytobenthos between the two study sites. **a**: Ratio of 16:0/16:1w7 (dimensionless). **b**: Chlorophyll *a* concentration (in µg.g sediment dry weight$^{-1}$). **c**: Relative contribution of various fatty acid classes . BRA = branched fatty acids, SFA = saturated fatty acids, MUFA = monounsaturated fatty acids and PUFA = polyunsaturated fatty acids

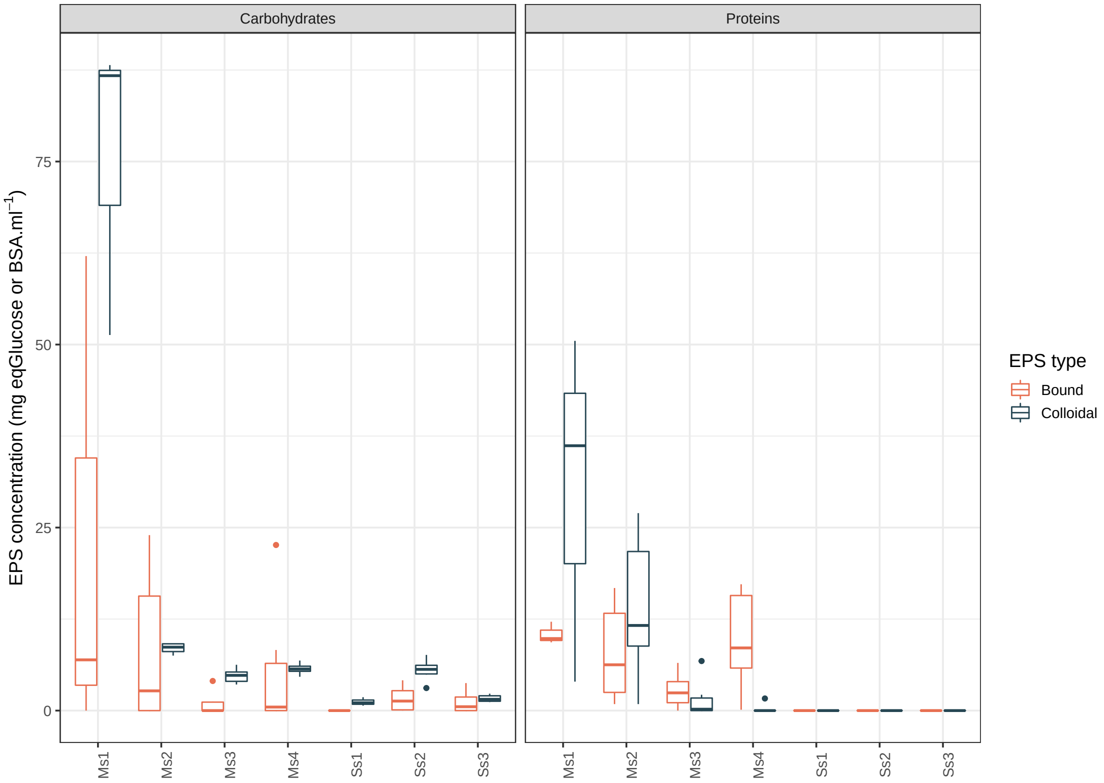


**Fig. SF2.** Colorimetric measurements of EPS concentrations in $\mathrm{mg}$ equivalent to Glucose or Bovine Serum Albumine (BSA) for carbohydrates and proteins respectively per $\mathrm{mL}$ of extracted EPS; colloidal EPS corresponded to loose, water-extractable exopolymers whereas bound EPS correspond to ion exchange resin-extractable exopolymers.

**Table ST1.** Presumed sources of colloidal and bound EPS (carbohydrates, proteins) at muddy and sandy sites. Position refers to the quality of alignment between fatty acids and EPS $\delta^{13}$C values: Aligned = the mean $\delta^{13}$C value of a given fatty acid was within the standard deviation or confidence interval of the corresponding EPS isotope ratio, Sup. (Superimposed)= $\delta^{13}$C values of EPS and fatty acids overlapped by their standard deviations or confidence intervals. **Bold underlined** = major fatty acid (20-40%) in the corresponding sources. **Bold** = important fatty acid (10-20%), *Italic* = present in trace amounts (<10%)

| Location | EPS type | Fatty acids | Position | Possible origin of FA |
|---|---|---|---|---|
| Mud | Colloidal | 22:6n-3 | Aligned | **Dinoflagellates**, **Haptophyta** (59), *Diatoms, Cyanobacteria* (24, 25, 59) |
| | | 20:0 | | *Bacteria* (55, 60) |
| | | 18:4n-3 | Sup. | **Haptophyta**, **Pheophyceae** (61), *Diatoms* (25) |
| | | 17:0iso | | **Bacteria** (55, 58) |
| | Bound | 20:4n-6 | Aligned | *Diatoms* (24, 25, 59), *Chlorophyta* (59) |
| | | 17:1n-5/7 | | *Bacteria* (60) |
| | | 16:4n-1 | | **Diatoms** (26), *Diatoms* (25, 59) |
| | | 22:0 | Sup. | *Diatoms* (26), *Cyanobacteria* (62) |
| | | 20:5n-3 | | **Diatoms** (24–26), **Diatoms** (59) |
| | | 18:4n-6 | | *Cyanobacteria* (63) |
| | | 18:2n-6 | | **Cyanobacteria, Chlorophyta** (59, 62), **Fungi** (64) |
| | | 18:1n-9/18:3n-3 | | **Cyanobacteria, Chlorophyta** (59, 62) / (24, 59) * |
| | | 17:0 | | *Bacteria, Diatoms* (26, 55) |
| | | 17:iso | | **Bacteria** (55, 58) |
| | | 16:3n-4/16:4n-3 | | *Diatoms* (59, 65)/ **Chlorophyta** (59) |
| | | 15:0 | | **Bacteria** (60), *Diatoms, Chlorophyta* (24, 25, 62) |
| Sand | Colloidal | 22:6n-3 | Aligned | **Dinoflagellates**, **Haptophyta** (59), *Diatoms, Cyanobacteria* (24, 25, 59) |
| | | 20:5n-3 | | **Diatoms** (26) |
| | | 20:4n-6 | | *Diatoms* (24, 25, 59), *Chlorophyta* (59) |
| | | 18:4n-6 | | *Cyanobacteria* (63) |
| | | 18:2n-6 | | **Cyanobacteria, Chlorophyta** (59, 62) |
| | | 18:1n-9/18:3n-3 | | **Cyanobacteria, Chlorophyta** (59, 62) / (24, 59) * |
| | | 18:1n-7 | | **Bacteria** (3) |
| | | 17:1n-5/7 | | *Bacteria* (60) |
| | | 17:0 | | *Bacteria, Diatoms* (26, 55) |
| | | 16:3n-4/16:4n-3 | | *Diatoms* (59, 65)/ **Chlorophyta** (59) |
| | | 16:1n-7 | | **Diatoms** (25, 26), **Cyanobacteria, Bacteria** (24, 55, 59, 62), *Chlorophyta* (62) |
| | | 16:1n-5 | | **Diatoms, Bacteria** (25, 26, 55) |
| | | 16:0 | | Major or important fatty acid in various sources (24–26, 55, 59, 61, 62, 66) |
| | | 15:0 | | **Bacteria** (60), *Diatoms, Chlorophyta* |
| | | 14:0 | Sup. | **Diatoms** (26) |
| | | 18:0 | | **Dinoflagellates** (59) * |
| | Bound | 15:0iso, 15:0anteiso | Aligned | **Bacteria** (57), *Bacteria* (3, 48, 55) |
| | | 18:1n-7 | Sup. | **Bacteria** (3, 28, 48, 55, 56), *Cyanobacteria, Chlorophyta, Diatoms* (24, 25, 59) |
| | | 18:0 | | **Dinoflagellates** (59) * |
| | | 16:0 | | Major or important fatty acid in various sources (24–26, 55, 59, 61, 62, 66) |
| | | 15:0 | | **Bacteria** (60), *Diatoms, Chlorophyta* |
| | | 14:0 | | **Diatoms** (26) |

* also detected in all sources in trace amounts (24–26, 55, 59–62, 64).
